# Tree diversity increases forest temperature buffering

**DOI:** 10.1101/2023.09.11.556807

**Authors:** Florian Schnabel, Rémy Beugnon, Bo Yang, Ronny Richter, Nico Eisenhauer, Yuanyuan Huang, Xiaojuan Liu, Christian Wirth, Simone Cesarz, Andreas Fichtner, Maria D. Perles-Garcia, Georg J. A. Hähn, Werner Härdtle, Matthias Kunz, Nadia C. Castro Izaguirre, Pascal A. Niklaus, Goddert von Oheimb, Bernhard Schmid, Stefan Trogisch, Manfred Wendisch, Keping Ma, Helge Bruelheide

## Abstract

Global warming is increasing the frequency and intensity of climate extremes. Forests may buffer such extreme events by creating their own microclimate below their canopy via cooling hot and insulating against cold macroclimate air temperatures. This buffering capacity of forests may be increased by tree diversity and may itself maintain forest functioning and biodiversity. However, despite its relevance for many ecosystem processes, the effect of tree diversity on temperature buffering is largely unexplored. Here, we show that tree species richness consistently increases forest temperature buffering across daily, monthly, and annual scales over six years. This finding is based on data from a large-scale tree diversity experiment covering a species richness gradient of 1 to 24 tree species. We found that species richness strengthened both components of forest temperature buffering: the attenuation of hot and of cold macroclimate air temperatures, with the cooling effect being more pronounced. The buffering effect of tree species richness was mediated by canopy density and structural diversity, assessed as leaf area index and stand structural complexity index, respectively. Safeguarding and planting diverse forests may thus mitigate negative effects of global warming and climate extremes on ecosystem functions and communities below the tree canopy.

## Introduction

Global warming and its impacts on the world’s forests^1^ are largely studied as effects of air temperatures measured outside forests in open-ground conditions (also referred to as macroclimate)^2^. However, this omits that forests can buffer temperature extremes such as hot and cold spells to some extent by creating their own microclimate below their canopy^3–5^, from which other organisms benefit, including sub-canopy trees. Among earth’s terrestrial ecosystems, forests are likely the one with the strongest air temperature buffering (hereafter ‘temperature buffering’) capacity owing to their often multi-layered canopies, which provide evapotranspirative cooling and shading, and decrease the mixing of air layers^5,6^. Temperature buffering occurs when microclimate temperature fluctuations are smaller than fluctuations in macroclimate temperatures^2^. Smaller temperature fluctuations below the canopy can be quantified as a lower temporal variance of temperatures, to which in the following we refer to as microclimate temporal stability^7^. The differences between macroclimate (outside forest) and microclimate (inside forest) temperatures are substantial, with global averages of –4.1 ± 0.5°C decreased temperature maxima and 1.1 ± 0.2°C increased temperature minima below the forest canopy^2^. This difference is larger than the average warming of land and ocean temperatures in 2001–2020 compared with 1850–1900 (0.8 to 1.1) °C^8^.

The temperature buffering capacity of forests has important consequences for forest functioning and biodiversity above- and belowground, especially in the context of global warming^2,4,9,10^. For instance, many physiological processes, such as photosynthesis or soil respiration^11^, scale exponentially with temperature, which implies that even small temperature increases may have large effects on rates, underlining the importance of temperature buffering. Furthermore, temperature buffering can influence forest biodiversity by slowing shifts in forest community composition towards warm-affinity species (i.e. thermophilization) under global warming^6,12,13^. However, the reciprocal control of tree diversity on forest temperature buffering remains largely unexplored.

Simulation studies showed that plant diversity can stabilise climate–vegetation feedbacks^14^. Moreover, tree species diversity has been shown to increase tree growth in mixtures^15–17^ and to enhance canopy complexity^18,19^, resulting in a greater thickness, density and structural diversity of the canopy layer (i.e., the buffering layer). It is thus conceivable that tree species richness may increase the temperature buffering capacity of forests by affecting these forest properties. For instance, mean tree height and the area of foliage per unit ground area (i.e. leaf area index; LAI)^20^ as proxies for the thickness and density of the buffering layer, modify the energy exchange at the canopy by influencing the penetration of sunlight and its albedo and evapotranspiration, which in turn affects the temperature buffering capacity of the forest^2^. Moreover, structural diversity^21–23^ measured, for instance, as stand structural complexity index (SSCI) from terrestrial laser scans^24^ may reduce the vertical mixing of air masses^25^ and thereby increase temperature buffering. However, there is very little empirical evidence for tree diversity effects on forest temperature buffering in general, and, in particular, regarding the mechanisms mediating such diversity–microclimate relationships. Establishing such causal relationships requires studies which experimentally manipulate tree species richness, and control for confounding factors, such as environmental variation or species identity effects^26,27^. A pioneering experimental study recently reported that tree species richness (1- vs 4-species) increased temperature buffering for some species mixtures^28^, but longer diversity gradients and data from multiple years would be necessary to generalise beyond specific species compositions and macroclimatic conditions as well as to understand the mediators of tree diversity effects on temperature buffering and their temporal dynamics.

Tree diversity effects on microclimate temperatures in forests may change over days, months and years. Compared with open-ground conditions, temperatures within forests are expected to be higher during night-time and winter, and lower during day-time and summer^10^. The underlying reason is that the energy exchange is shifted from the ground surface to the canopy^29^. Consequently, forest canopies mitigate hot temperatures via evapotranspiration (consumption of latent heat), reflecting or absorbing solar radiation and emitting long-wave radiation, and insulate against cold temperatures via heat retention^3,5^. However, many more processes may be involved depending on the spatiotemporal scale studied^2^. For example, evapotranspirative cooling effects decrease with decreasing soil water availability^2^, highlighting the potential influence of inter-annual dynamics and extremes in macroclimatic conditions (such as droughts) for temperature buffering. However, the relative importance of tree diversity effects on temperature buffering across temporal scales remains unknown.

Here, we analyse microclimate measurements conducted within forests of 1 to 24 tree species covering six years (2015–2020) from a large-scale subtropical tree diversity experiment (BEF-China^15,26^). Assembling the communities with varying species richness randomly from species pools resulted in stands that differ in canopy thickness^15^, density^30^ and structural diversity^19^. We aim to understand the role of tree species richness and these mediating factors for temperature buffering below forest canopies at different temporal scales (i.e. daily, monthly and yearly). In our subtropical study system, which is characterised by a monsoon climate, high macroclimate temperatures coincide with high water availability for evapotranspirative cooling, which should promote temperature offsets between micro- and macroclimate, particularly for maximum temperatures^2^. Hence, we expect tree species richness effects on microclimate to be most pronounced for the buffering of maximum temperatures. We tested the following hypotheses: H1: tree species richness increases the temperature buffering potential of forest canopies via cooling hot and insulating against cold macroclimate temperatures at daily, monthly, and annual time scales. H2: species richness effects on temperature buffering— measured as microclimate temperature stability—are consistently positive across time scales but strongest when macroclimate temperatures are high. H3: positive tree species richness effects on temperature buffering are mediated by enhanced canopy thickness, density, and structural diversity.

## Results

On the daily scale, we found below-canopy air temperatures to decrease with tree species richness during daytime, while they increased with species richness during the night (Fig. 1a). Hence, the mode of tree species richness effects on microclimate temperature changed significantly with the diurnal course in macroclimate temperatures from positive (during cold night-time hours) to negative (during hot day-time hours; Fig. 1a: interaction between species richness and hour significant at P < 0.001). Mitigating species richness effects on microclimate temperature were strongest at midday peak hours (mean temperature offsets of –2.5 ± 0.2°C from noon to 3 pm) and positive effects strongest around midnight (+0.4 ± 0.04°C from 11 pm to 2 am) between stands with 1 and 24 tree species, respectively.

**Figure 1.**
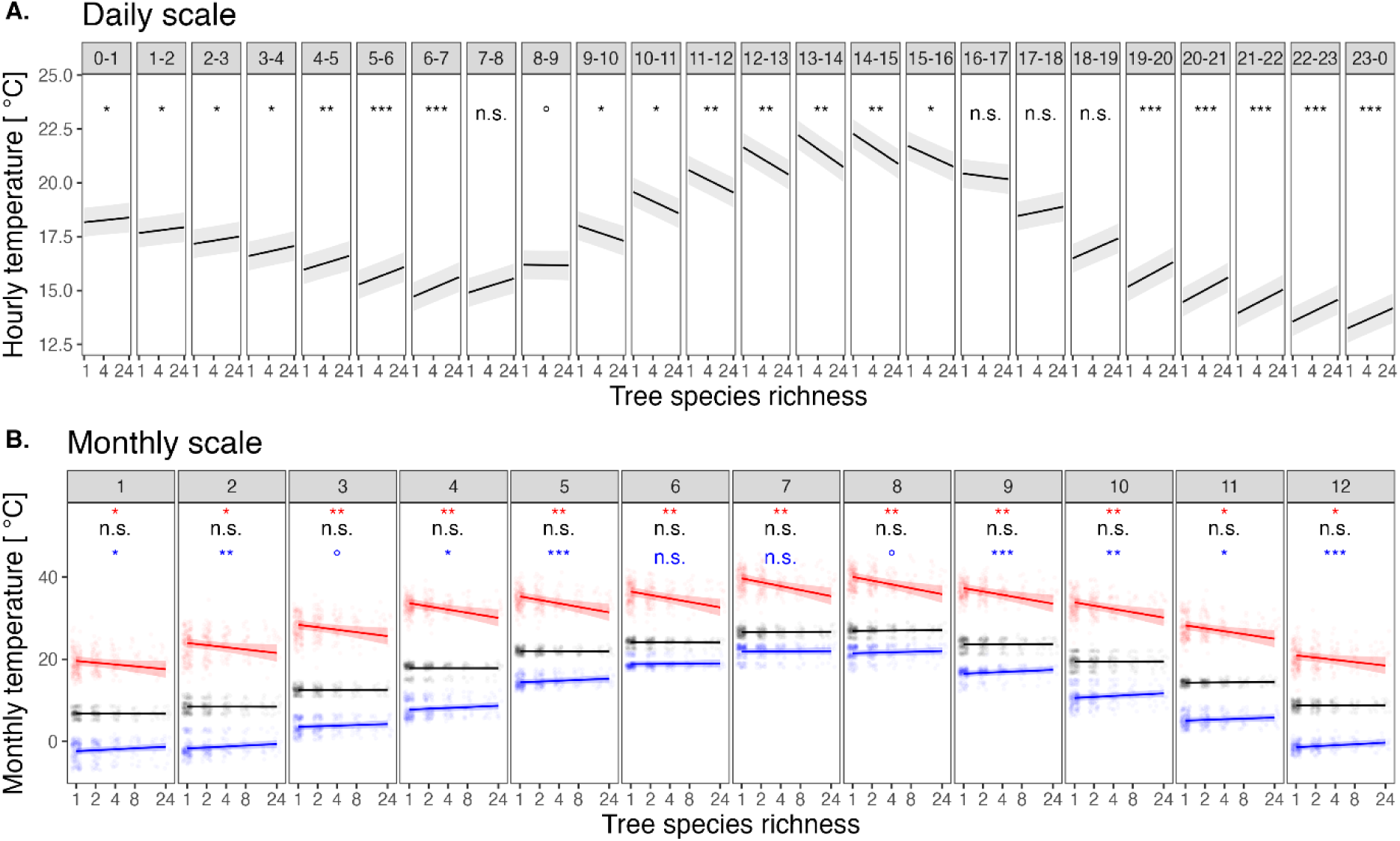
Tree species richness effects on microclimate temperature on (A) the daily and (B) the monthly scale. (A) Hourly modulation of microclimate temperatures by tree species richness (n = 63 plots and 4 million values). (B) Monthly modulation of maximum (red), median (black) and minimum (blue) daily temperatures per month by tree species richness (n = 63 plots and 4476 values). Lines show predictions of linear mixed-effects models, and shaded bands indicate 95% confidence intervals. Data points in (B) are jittered to enhance visibility. Species richness was log-transformed in all models. See Supplement S2 for complete model outputs. Significance levels: “n.s.”: non-significant, “°”: p<0.1, “*”: p<0.05, “**”: p<0.01, and “***”: p<0.001.

On the monthly scale, we examined maximum, minimum and median daily microclimate temperatures across months (Fig. 1b). We found that tree species richness significantly reduced maximum microclimate temperature across all months (January–December, P-value range of slopes 0.002–0.033); this buffering effect was strongest during summer (up to –4.4°C ± 0.6°C in 24-species mixtures in July, P = 0.004) and during high macroclimate temperatures (Supplement S1). Tree species richness also increased minimum microclimate temperatures in most months (September–May; P-value range of slopes 0.001–0.053); this buffering effect was strongest in winter (up to +1.1°C ± 0.2°C in 24-species mixtures in December, P < 0.001), non-significant during summer (June–August; P > 0.05), and strongest during low macroclimate temperatures (Supplement S1). We found no significant effect of tree species richness on median monthly temperatures (Fig. 1b; P > 0.5 for all months), i.e., species richness only affected temperature extremes. Hence, as hypothesised, tree species richness cooled hot and insulated against cold macroclimate temperatures, which contributed to enhanced buffering of temperature extremes in species-rich stands.

We quantified the temperature buffering capacity of a tree community on monthly and annual time scales as the temporal stability^7^ of microclimate temperature, calculated as the inverse of the coefficient of variation (CV) of hourly temperature measurements. This stability metric is commonly used in biodiversity–ecosystem functioning studies to provide insights into the stabilising effects of biodiversity for multiple ecosystem processes and at different levels of organization^31–33^. We found a consistently positive effect of tree species richness on monthly temperature buffering across the entire year (January–December; P ≤ 0.006 for all months), which was strongest in summer (June–August; Fig. 2a) and during high macroclimate temperatures (Supplement S1). Tree species richness also had significant positive effects on annual temperature buffering during all years examined (P < 0.001; Fig. 2b). Effects of species richness on annual temperature buffering (slope of the species richness–temperature buffering relationship) did not change significantly across years, but the absolute temperature buffering capacity of the examined forest communities changed with macroclimatic conditions. Temperature buffering was significantly lowest during the driest year (i.e. 2018, the year with the lowest SPEI values, P < 0.001; Fig. 2b, Supplement S2).

**Figure 2.**
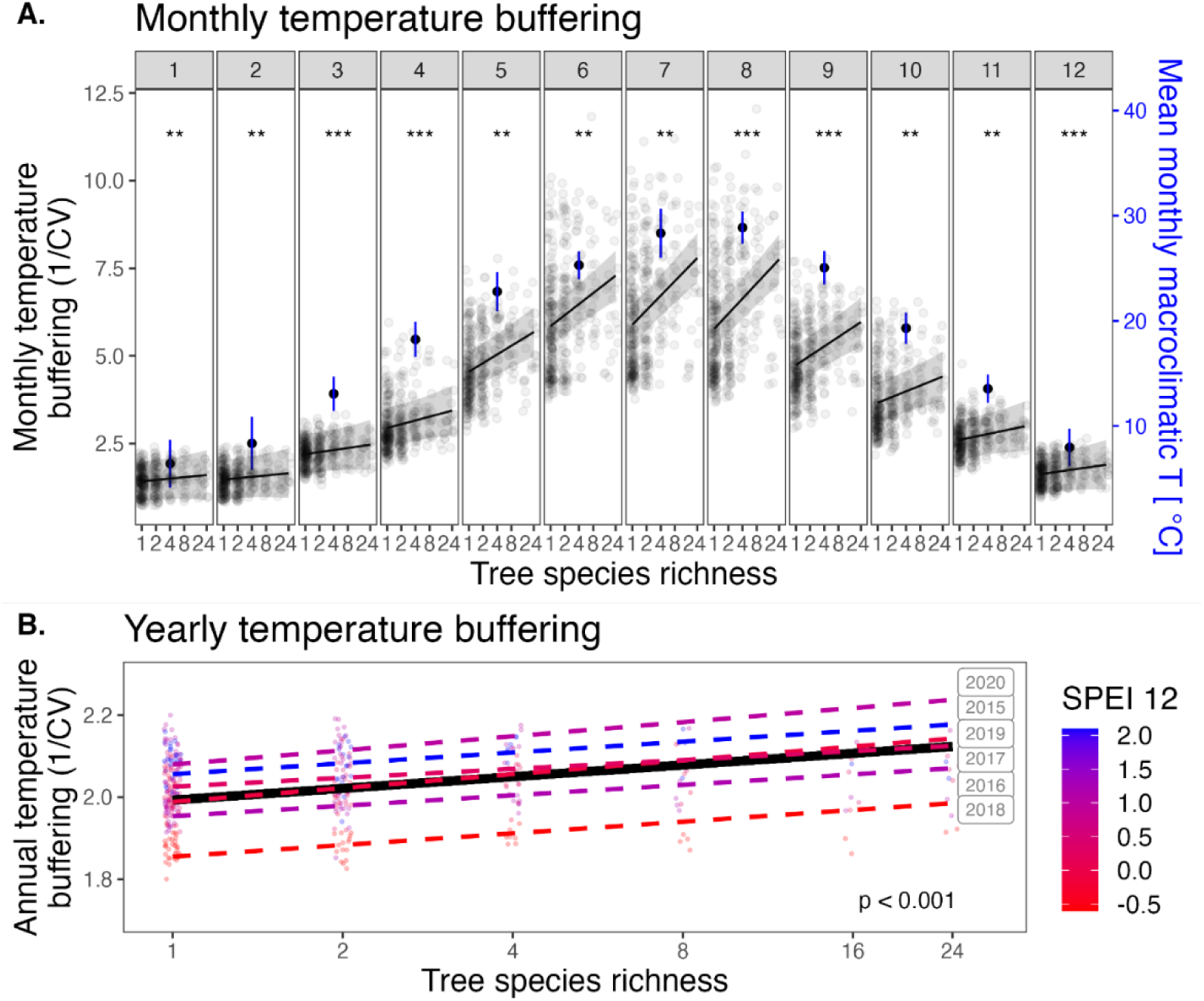
Tree species richness effects on temperature buffering on (A) the monthly and (B) the annual scale. (A) Modulation of monthly microclimate temperature stability (n = 63 plots and 4476 values) by tree species richness and month of the year. (B) Modulation of annual microclimate temperature stability (n = 63 plots and 375 values) by tree species richness and year. In all panels, the lines show predictions of linear mixed-effects models, and data points are jittered to enhance visibility. In (A), p-values refer to the effects of species richness on monthly temperature buffering and solid black points show mean monthly macroclimate temperatures, while shaded bands and blue whiskers indicate 95% confidence intervals. In (B), the P-value refers to the effect of species richness across years. Points and lines are coloured according to their value with deeper red and blue indicating increasing and decreasing drought, respectively, based on annual values of the standardised precipitation evapotranspiration index (SPEI12). Species richness was log-transformed in all models. See Supplement S2 for complete model outputs. Significance levels: “n.s.”: non-significant, “°”: p<0.1, “*”: p<0.05, “**”: p<0.01, and “***”: p<0.001.

We used piecewise Structural Equation Models (SEMs; Fig. 3) to examine potential mechanisms that may mediate the observed tree species richness effects on monthly temperature buffering (Fig. 2). Out of a set of potential variables and based on literature-derived hypotheses (Supplement S3), we selected mean tree height, LAI, and SSCI as measures of canopy thickness, density, and structural diversity, respectively. Once controlling for the effect of macroclimate, we found LAI to have the strongest positive effect on temperature buffering (Std. estimate = 0.33, P = 0.011), followed by SSCI (Std. estimate = 0.23, P = 0.002), while mean tree height had no significant effect on temperature buffering (P = 0.5, Fig. 3A). Both LAI and SSCI significantly increased with increasing tree species richness (Std. estimate = 0.82, P = 0.007 and Std. estimate = 0.17, P = 0.002, respectively). Once accounting for these forest properties and their influence on temperature buffering, we found no remaining direct effect of tree species richness on temperature buffering (P = 0.3, Fig. 3A). Using tree basal area measured in 2019 (another commonly used proxy for canopy- or stand density^28,34^) instead of LAI resulted in similar pathways: tree species richness increased basal area (Std. estimate = 0.23, P = 0.049), which in turn enhanced temperature buffering (Std. estimate = 0.41, P = 0.011; Supplement S4). The influence of the different drivers changed over the annual course (Fig. 3B): LAI was the strongest driver of temperature buffering during the growing season (March– September), while SSCI mostly affected temperature buffering before and after the growing season.

**Figure 3.**
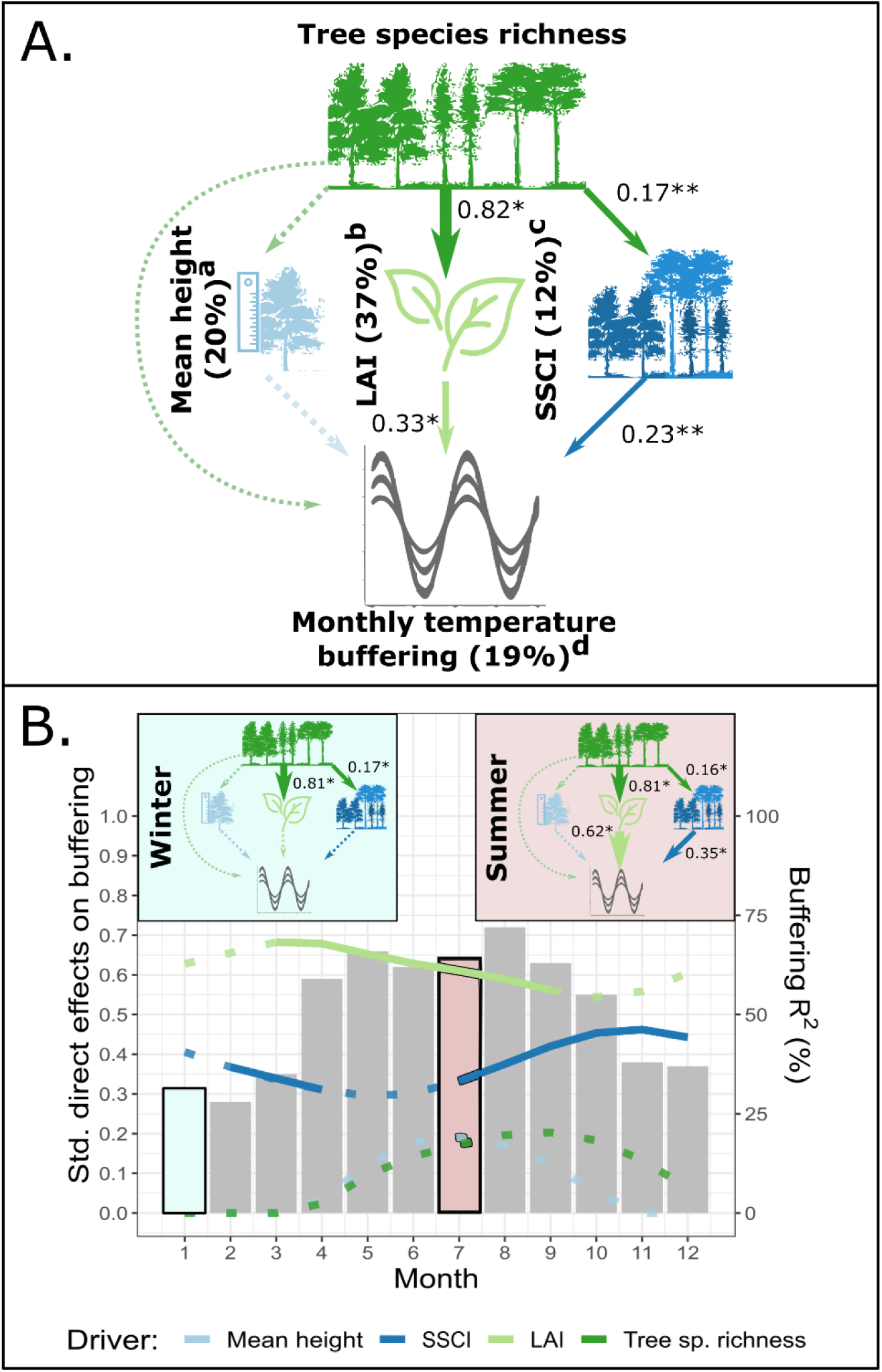
Structural Equation Models (SEMs) examining potential mediators of tree species richness effects on monthly temperature buffering. The SEMs test the direct effects of tree species richness and its indirect effects mediated by mean tree height, leaf area index (LAI), and stand structural complexity index (SSCI) on monthly temperature buffering while controlling for macroclimate temperatures. The SEM in (A) was fit to microclimate data of all months, and tree species richness effects on forest properties (i.e. mean tree height, LAI, and SSCI) were tested on the datasets built for this purpose (^a^: 32 plots, ^b^: 54 plots, ^c^: 74 plots, ^d^: 27 plots; see methods). All pathways were fit to data from site A measured in 2019 except for LAI, which was measured in 2014 (see methods and Supplement S3 for details). Species richness and SSCI were log-transformed in all models. Significant directional relationships between variables are shown as solid and nonsignificant relationships as dashed arrows. Significant standardised path coefficients are shown next to each path (*P < 0.05, **P < 0.01, and ***P < 0.001), and path width is scaled according to coefficient size. The explained variation of variables (marginal R^2^) is given in %. The SEM fit the data well (Fisher’s C = 8.4, df = 6, P = 0.21). In (B), the same SEM was fit separately for each month to explore temporal trends in the path coefficients. The SEMs in January and July exemplify pathways during winter and summer, respectively. For each month, coloured curves show standardized path coefficients (dashed if non-significant) and bars show the variation in temperature buffering explained by the examined forest properties (marginal R^2^); note that the marginal R^2^ in (A) is lower than in the monthly models in (B) as it only captures the variation explained by fixed effects, which do not account for the strong variation in temperature buffering between months. See Supplement S4 for complete model outputs.

## Discussion & Outlook

In a large-scale tree diversity experiment, we observed a consistent increase in forest temperature buffering across daily, monthly, and annual scales with increasing tree species richness. Confirming H1, species-rich forests cooled high and insulated against cold macroclimate temperatures better than species-poor forests. This positive effect had a considerable magnitude with –4.4°C (± 0.6°C) and +1.1°C (± 0.2°C) in peak summer and winter for monocultures *vs* 24-species mixtures, respectively. Confirming H2, temperature buffering was thus driven primarily by a reduction of maximum below-canopy temperatures, with this effect being strongest during hot macroclimate conditions (during midday and summer).

We expected species-rich tree canopies to mainly cool hot temperatures by enhancing evapotranspiration and the reflection of short-wave and emittance of long-wave radiation^3,5^. Likewise, tree canopies may insulate against cold temperatures by retaining heat and long-wave radiation, even though many more processes are likely involved^2^. Consistent with our findings, stronger buffering of maximum relative to minimum temperatures predominates across the world’s forests^2,3^. Moreover, next to temperature extremes, droughts will likely threaten the world’s forests during the 21^st^ century^1,35^. We found the lowest absolute temperature buffering in the driest year (2018) of our observation period (Fig. 2), likely due to reduced cooling potentials via evapotranspiration (as a result of the low atmospheric and soil moisture)^2^. However, the buffering role of tree species richness was maintained (Fig. 2), indicating that tree species richness provides an insurance against climate extremes in the subtropical tree communities we studied.

There is ample evidence that forests buffer temperature extremes^3–5^ and that species identities matter for temperature buffering^25,28^, but the role of tree diversity has largely remained hidden. The few former studies on the role of tree species composition for temperature buffering reported predominantly non-significant effects of species richness^25,34,36^. Positive effects were rare and only found for specific mixtures^28^. It may be that idiosyncrasies of the investigated species prevented the detection of general patterns of species richness in earlier studies or that the level of species richness analysed was too low to detect significant effects. Our experimental design with a long tree species richness gradient ranging from 1 to 24 tree species and randomised extinction scenarios where each richness level was represented by different species compositions and each species occurred at each richness level^15,26^ allowed us to move beyond the effects of specific species compositions while controlling for environmental variation and species identity effects. Confirming this view, earlier studies in our experiment have demonstrated that most species and not only some particular species contributed to the observed diversity effects. For instance, complementarity and not selection effects drove the net positive tree diversity effects on stand volume^15^. Moreover, at the examined scale, this high tree species richness level is common under natural conditions in the subtropical study region^26^.

Partially confirming H3, we found positive tree species richness effects on temperature buffering to be mediated by enhanced canopy density and structural diversity but not by canopy thickness (Fig. 3). The absence of a remaining direct tree species richness effect after accounting for these forest properties supports the use of the chosen proxies (LAI and SSCI) and suggests that we captured the dominant mechanisms driving temperature buffering. Still, monitoring other potential drivers, such as enhanced transpiration^37^, will be relevant for comprehensively understanding species richness effects on temperature buffering. Canopy density and structural diversity were already shown to be enhanced by tree species richness in our experiment^19,30^ and elsewhere^16,17,24^. Likewise, canopy density^2,34,38^ and structural diversity^24,25,36^ were reported to be significant drivers of forest temperature buffering. Moreover, and similar to our findings, structural diversity was more relevant than mere canopy height in this context^25^. However, these studies did not elucidate the mechanistic links between species richness, canopy density, structural diversity, and temperature buffering. Here, we provide experimental evidence that most species richness effects on temperature buffering are indirect and mediated via diversity-induced changes in these forest properties. This notion is consistent with canopy cover, another proxy for canopy density, mediating species richness effects on minimum and maximum temperatures^28^. Furthermore, our study reveals that drivers of temperature buffering in forests exhibit temporal complementarity, with LAI being most relevant during the peak growing season and SSCI, which captures the structural diversity of canopy elements (stems and branches) during the leaf-off period of the deciduous tree species, taking over outside the growing season.

The positive effect of tree diversity on temperature buffering we report represents, as also highlighted by advances in grassland research^39,40^, a previously overlooked biodiversity-ecosystem functioning (BEF) relationship, with potentially far-reaching implications. In contrast to other mechanisms that cause positive BEF relationships in forests, such as biotic interactions between trees, negative density effects, or multitrophic interactions^41^, which are all species-specific, temperature buffering emerges from the community as a whole. The resulting lower temperature variation in species-rich forests may safeguard ecosystem functions, particularly those that respond non-linearly to temperature^11^, against temperature maxima (and minima). This may be especially relevant for functions severely impeded beyond narrow threshold ranges of temperature, such as net photosynthesis rates^42^. Likewise, belowground functioning, including carbon sequestration, decomposition, and nutrient cycling^10,43,44^, may be enhanced by temperature buffering. As a result, trees in mixtures may grow^45,46^ and regenerate better^47^ in ameliorated microclimates, which may, in turn, enhance temperature buffering via enhancing canopy density (Fig. 3). Moreover, by reducing maximum temperatures (Fig. 1), tree diversity-enhanced temperature buffering may impact forest biodiversity under global warming by reducing the thermophilization of below-canopy communities^6,12,13^. Finally, forest temperature buffering also alleviates heat stress for humans, and our findings indicate that tree species richness may amplify this effect far stronger than previously reported^34^.

Overall, we suggest that preserving and planting diverse forests^48^ is a promising approach to increase the temperature buffering function of forests, thereby protecting ecosystem functions and communities below the tree canopy against global warming. We compared the effects of increasing tree diversity on temperature buffering and the mediation of tree diversity effects by LAI and SSCI at constant planting density. Hence, at higher planting densities, mixtures would still outperform monocultures. Nonetheless, attempting to promote LAI and, thereby, temperature buffering by planting monocultures with only a single or a few shade-tolerant tree species may be theoretically possible. However, such species-poor forests would have other well-known limitations, such as a higher susceptibility to specialist pests and pathogens, droughts, and storms^48^. In contrast, species-rich forests are more likely to maintain their buffering capacity in the future^28^, given their higher stability under global change^31^, while simultaneously providing a broader range of ecosystem services^48^. Despite examining young planted forests (up to 11 years after establishment), we already detected a strong temperature buffering capacity. Our findings thus highlight the benefits of diverse planted forests for large-scale forest restoration initiatives^49^ and urban forests that aim at reducing thermal stress in a warming world.

## Methods

### Study site and experimental design

We used data from a large-scale tree biodiversity experiment, the Biodiversity–Ecosystem Functioning China Experiment (BEF-China experiment)^15,26^, located in Xingangshan, Dexing, Jiangxi (29°08′–29°11′ N, 117°90′–117°93′ E). The experiment was established at two sites, A and B, which were planted in 2009 and 2010, respectively. Each site covers approximately 20 ha in size. The site’s climate is governed by the subtropical monsoon, with cold and dry winters and hot and humid summers. The mean annual temperature and precipitation are 16.7°C and 1821 mm (mean from 1971–2000)^50^. Inter-annual changes in climate-induced water availability are strong and driven primarily by changes in precipitation and only to a lower degree by changes in temperature^15,31^. The native forests of the study region harbour a high tree species richness and are dominated by broadleaf tree species^26^. Based on a total pool of 40 native evergreen and deciduous broadleaf tree species, we created manipulated species richness gradients of 1 to 24 coexisting species (Supplement S5). Overall, 226,400 individual trees were planted on 566 plots, with each plot featuring a size of 25.8 × 25.8 m^2^ (1/15 ha) and 400 trees. To increase generality and statistical power, tree species were allocated to different extinction scenarios following a broken-stick design with partly overlapping species pools per extinction scenario^15,26^. Here, we used data from extinction scenarios to which species were randomly allocated, specifically the 64 Very Intensively Studied Plots (VIPs) of the BEF-China experiment^26^.

### Micro- and macroclimate measurements

The microclimate air temperature was recorded hourly over six years (January 2015–December 2020) across the VIP plots (32 at each site) using temperature loggers (HOBO Pro v2, U23-001) covered by a rain-protection shield and installed at 1 m height in the centre of the plots (see Supplement S5). Data were controlled and cleaned to remove unrealistic data due to logger malfunction (e.g., temperature outliers or time series divergent dynamics; Supplement S5). Plots with incomplete monthly records were excluded from the monthly analyses and incomplete yearly records were excluded from yearly analyses (1 plot of the 64 plots was removed in all analyses; Supplement S5). Macroclimate data—minimum, average, and maximum monthly temperature (°C), monthly precipitation sum (mm) and monthly potential evapotranspiration (mm) sum—were retrieved from the high-resolution gridded dataset of the Climatic Research Unit (CRU) Time-Series (TS) version 4.06^51^ with a 0.5° (latitude/longitude) resolution, which is based on interpolated climate station observations. To explore if diversity– microclimate relationships were influenced by water availability, we further calculated the Standardised Precipitation-Evapotranspiration Index (SPEI)^52^ based on these precipitation and evapotranspiration data with the SPEI package^53^. The SPEI is a commonly used drought index that captures the climatic water balance (precipitation minus potential evapotranspiration) at different time lengths from a single month (SPEI1) to an entire year (SPEI12; January– December). SPEIs below –1 and above 1 can be considered exceptionally dry or wet compared to the average conditions during a climate reference period^54^ (here 1901–2019).

### Temperature buffering and stability

Using the hourly microclimate temperature measurements, we calculated different measures describing temperature extremes and temperature buffering. We calculated monthly minimum, median and maximum microclimate temperatures per plot. Minimum and maximum monthly temperatures were calculated by taking the median of the 5% lowest and 95% highest temperatures, respectively. We quantified temperature buffering on monthly and annual time scales as the temporal stability (*S*)^7^ of microclimate temperature, calculated as:

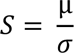

Where µ and σ are the mean and standard deviation of hourly temperature measurements per month or year, hereafter referred to as monthly or annual temperature stability.

### Assessment of microclimate drivers

We assembled a range of variables describing canopy thickness, density, and structural diversity from former studies and tree inventories in the BEF-China experiment. Out of these potential variables, we selected the ones with the highest relevance for temperature buffering according to literature-derived hypotheses (focussing on the ones most successfully used as predictors of temperature buffering in other studies; Supplement S3), and compared correlations between variables (Supplement S4). Specifically, we selected mean tree height, leaf area index (LAI), and Stand Structural Complexity Index (SSCI) to describe canopy thickness, density, and structural diversity, respectively. Tree height was measured as the mean height of the central 6 × 6 trees in each plot to avoid edge effects, as described in Huang et al.^15^. LAI was measured using digital hemispheric photography at five positions within each plot in August by Peng et al.^30^, and SSCI by a single terrestrial laser scan at the centre of each plot under leaf-off conditions of the deciduous tree species (February–March) as described by Perles-Garcia et al.^19^ (see these studies for details on the respective methods and datasets). For all forest property variables, we used data collected at site A of the BEF-China experiment in 2019 (where we had the best data coverage), except for LAI, measured in 2014.

### Statistical analyses

We used linear mixed-effects models (LMMs) to test for the effects of tree species richness on microclimate temperatures and temperature buffering across time scales and VIP plots (n = 63 plots, tree species richness ranging from 1–24 species). We tested for species richness effects on hourly temperatures and on minimum, median and maximum monthly temperatures using LMMs in which species richness in interaction with hour or month were considered fixed effects (Supplement S2). Similarly, we tested for species richness effects on monthly and annual temperature stability using LMMs in which species richness in interaction with month or calendar year were considered fixed effects (Supplement S2). We accounted for the experimental design of our study through a nested random effect structure of plots nested within the experimental site (A or B) and for temporal autocorrelation by using a first order autocorrelation structure (corCAR1) for time covariates (days, months or years). Additionally, we explored how diversity effects, i.e. the slopes of the regressions between species richness and monthly minimum, median and maximum microclimate temperatures and monthly temperature buffering (Figs. 1–2), depended on macroclimate conditions (monthly minimum, average and maximum temperatures and SPEI values) (Supplement S1). At the annual scale, we tested if temperature stability was related to annual climatic water balances by replacing calendar years by annual SPEI values in the respective LMM (Supplement S2).

To examine the mechanisms that may mediate tree species richness effects on temperature buffering, we used Structural Equation Models (SEMs). The hypothesis-driven SEMs were informed by previous work, including from the herein-examined experiment (see Supplement S3 for the conceptual model and the literature-derived hypotheses). Specifically, we examined if canopy thickness, density, and structural diversity, captured by mean tree height, LAI and SSCI, respectively, mediate tree species richness effects on temperature buffering. We accounted for potential correlations between these forest properties through including partial, non-directional correlations between them. We controlled for monthly variations in macroclimate temperatures by dividing monthly temperature buffering values through monthly macroclimate temperature values. Species richness effects on temperature buffering changed over the year (Fig. 2a). Hence, to capture potential temporal changes in the strength of the examined drivers, we fit separated SEMs for each month. In contrast, the strength of species richness effects on temperature buffering did not change significantly between years (Fig. 2b), which allowed us to focus on the year for which we had the most data on forest properties. We thus explored how species richness affected temperature buffering via canopy thickness, density and structural diversity in 2019 at site A, where we had measurements of all forest properties (except for LAI which was measured in 2014) and where temperature buffering was close to the mean response across years (Fig. 2b). Using stand-level basal area measured in 2019 instead of LAI resulted in similar pathways (Supplement S4). To remain consistent with prior knowledge on relationships between species richness and the examined forest properties in our experiment, we fitted direct pathways between species richness and LAI and SSCI using the datasets and model structures from the original studies (refs^19,30^). Therefore, the tree species richness–forest properties models were fitted on larger plot sets (n = 32, 54, and 74 plots for mean tree height, LAI, and SSCI, respectively) than the forest properties–temperature buffering models fit for the plots for which we had microclimate data and data on all examined forest properties (n = 27 plots, tree species richness ranging from 1–16 species). To prevent pseudo-replication caused by measuring tree height, LAI and SSCI on an annual basis, relationships between tree species richness and these forest properties were fitted using yearly datasets instead of monthly ones. In the tree species richness–LAI model, we included terms correcting for very large residual effects resulting from the presence of few specific species in the examined tree communities following Schmid et al. 2017^55^ as detailed in Peng et al.^30^. We assessed global model fit via Fisher’s C statistic (P > 0.05) and the independence of variables with tests of direct separation (P < 0.05 for violation of independence) and posteriori, included partial, non-directional correlations between non-independent variables^56^ (Supplement S4).

All data handling and statistical analyses were performed using the R statistical software version 4.1.3. Explanatory variables in the SEMs were centred and divided by one standard deviation using the ‘scale’ function, to avoid any model-fit deviation due to scale differences between variables. Tree species richness was log2-transformed in all models. LMMs and individual SEM pathways were fit with the nlme package^57^ and SEMs with the piecewiseSEM package^56^. Model assumptions (i.e. normality, independence and homogeneity of variance, and independence of explanatory variables) were tested with the ‘check_model’ function in the performance package^58^.

## Data availability

The datasets generated and analysed in the study will be made publicly available upon publication via the BEF-China project repository, at http://data.botanik.uni-halle.de/bef-china

## Code availability

All R scripts used for this study can be found in our GitHub repository, at https://github.com/remybeugnon/Schnabel-Beugnon-Yang-et-al_tree-diversity-temperature-buffering

## Acknowledgements

We thank local workers for their help in the field. This research was supported by the Deutsche Forschungsgemeinschaft (DFG, German Research Foundation; grant DFG FOR 891), the International Research Training Group TreeDì jointly funded by the DFG (grant 319936945/GRK2324) and the University of Chinese Academy of Sciences (UCAS). We are grateful for the support of iDiv funded by the DFG (grant DFG-FZT 118, 202548816). F.S. acknowledges support by a TreeDì start-up grant. R.B. acknowledges funding by the Saxon State Ministry for Science, Culture and Tourism (SMWK; grant 3-7304/35/6-2021/48880), N.E. funding by the DFG (grant Ei 862/29-1), H.Y. and N.E. funding by the DFG (grant FOR 5000), and B.S. support by the University Research Priority Program Global Change and Biodiversity of the University of Zurich.

## Author Contributions

H.B., K.M., B.Y., W.H., P.A.N, G.v.O., B.S. and C.W. designed the experiment; F.S., R.B. and R.R. conceived the study; B.Y., H.B., F.S., R.B., X.L., A.F., M.D.P.G., G.H., W.H., M.K., N.C.C.I., P.A.N., G.v.O. and S.T. measured and/or compiled data; F.S., R.B., R.R., B.Y., N.E., Y.H., X.L., C.W., and H.B. developed and refined the analysis concept; R.B. analysed the data with support by F.S. and G.H; F.S., R.B., R.R., B.Y., S.C., N.E., Y.H., C.W. and H.B interpreted the data; R.B. created figures; F.S. wrote the manuscript with support by R.B; F.S., R.B., B.Y., R.R., N.E., Y.H., C.W., S.C., A.F., M.D.P.G., G.H., W.H., M.K., X.L., N.C.C.I., P.A.N., G.v.O., B.S., S.T., M.W., K.M. and H.B. contributed substantially to revisions of drafts.

The authors declare no competing interests

Supplementary Information is available for this paper at: Suppl.S1–S5

