## Supplementary Material for "Tree diversity increases forest temperature buffering"

#### Contents

Monthly diversity effects *vs.* macroclimate . . . . . . . . . . . . . . . . . . . . . . . . . . . . . . . . 2

#### Monthly diversity effects *vs.* macroclimate

##### Maximum temperature

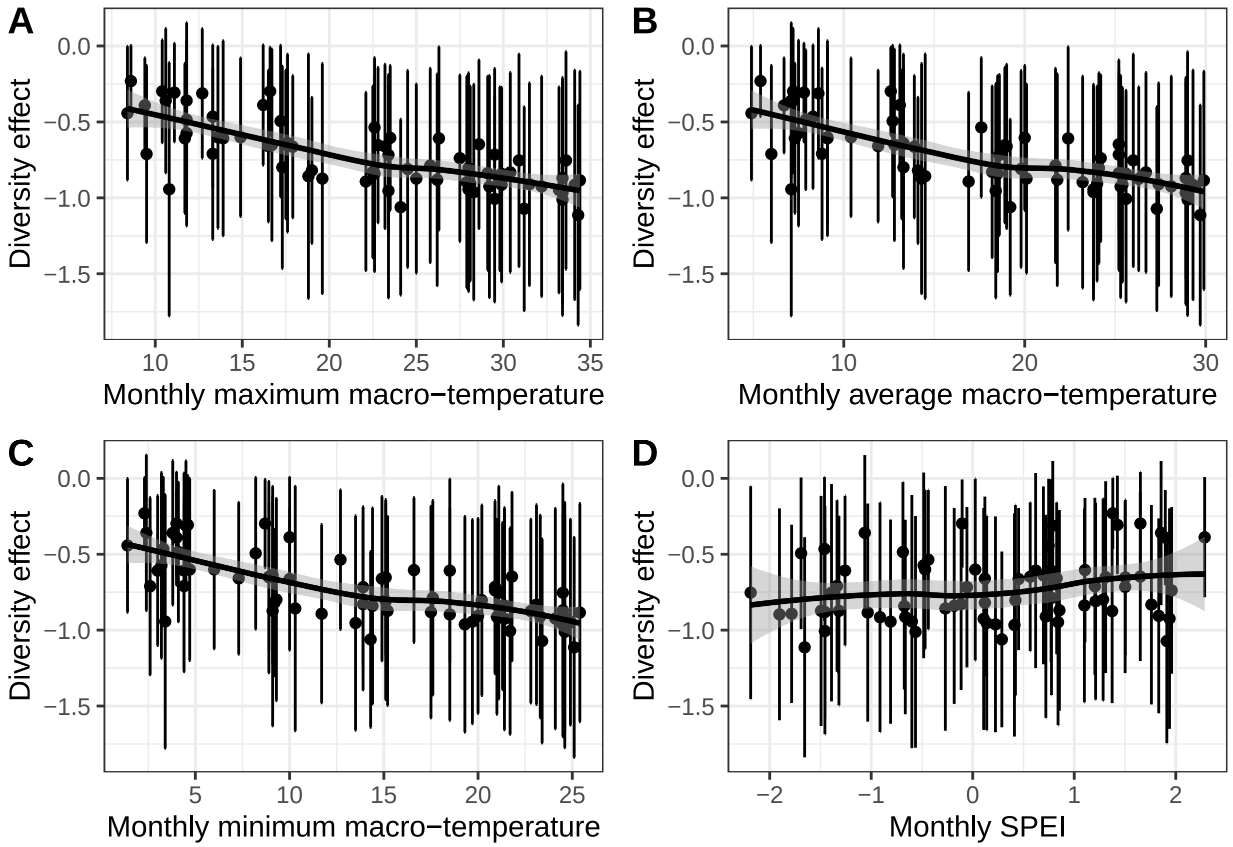

Figure S1 Macroclimate effects on the relationship between tree species richness and maximum microclimate temperature. Black points and error bars show diversity effects, i.e., slopes and respective standard errors, of the regression between maximum microclimate temperatures and tree species richness per month based on the model in Fig.1b (n = 63 for 12 months and 6 years). The change in these diversity effects with macroclimate conditions is examined for maximum (A), average (B) and minimum (C) monthly macroclimate temperatures and monthly values of the standardised precipitation evapotranspiration index (SPEI1; D). Trends are highlighted with a loess regression.

**Median temperature**

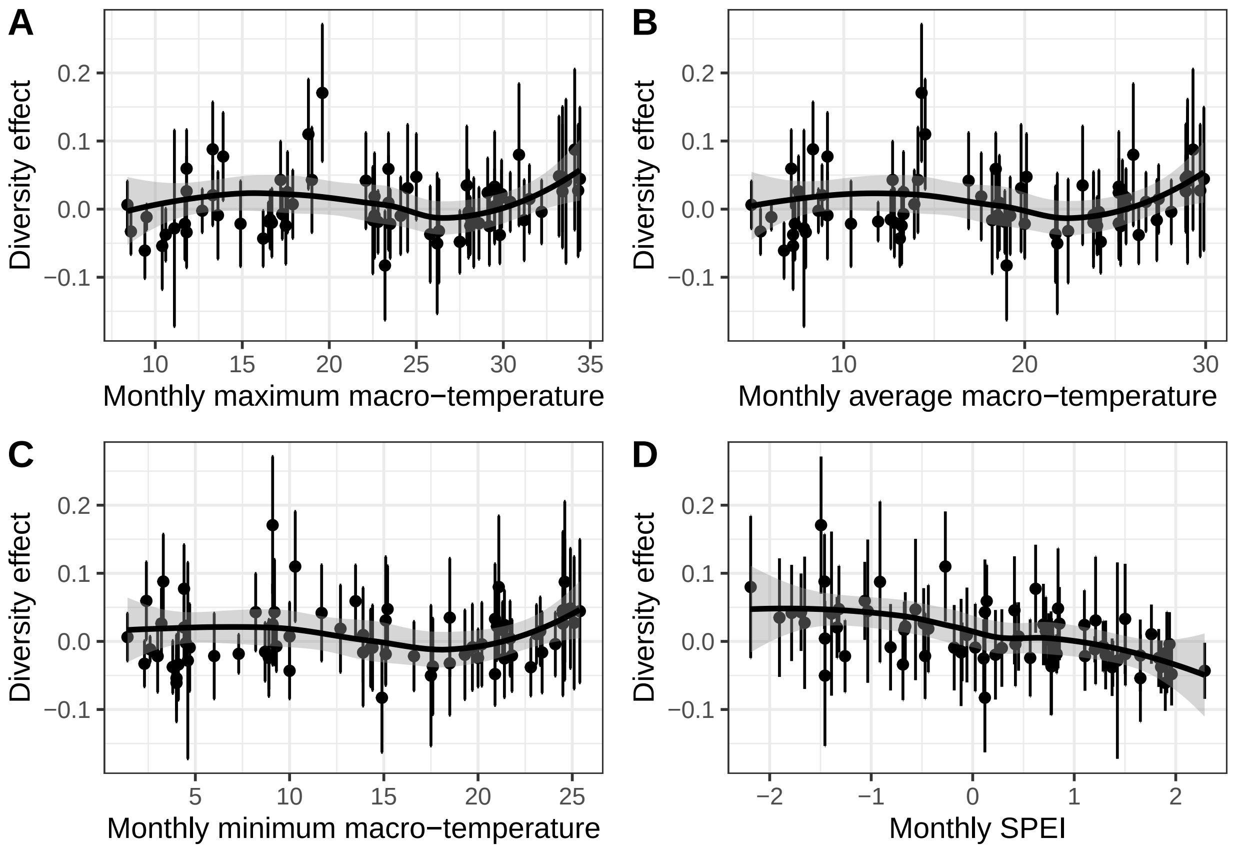

Figure S2 Macroclimate effects on the relationship between tree species richness and median microclimate temperature. Black points and error bars show diversity effects, i.e., slopes and respective standard errors, of the regression between median microclimate temperatures and tree species richness per month based on the model in Fig.1b (n = 63 for 12 months and 6 years). The change in these diversity effects with macroclimate conditions is examined for maximum (A), average (B) and minimum (C) monthly macroclimate temperatures and monthly values of the standardised precipitation evapotranspiration index (SPEI1; D). Trends are highlighted with a loess regression.

**Minimum temperature**

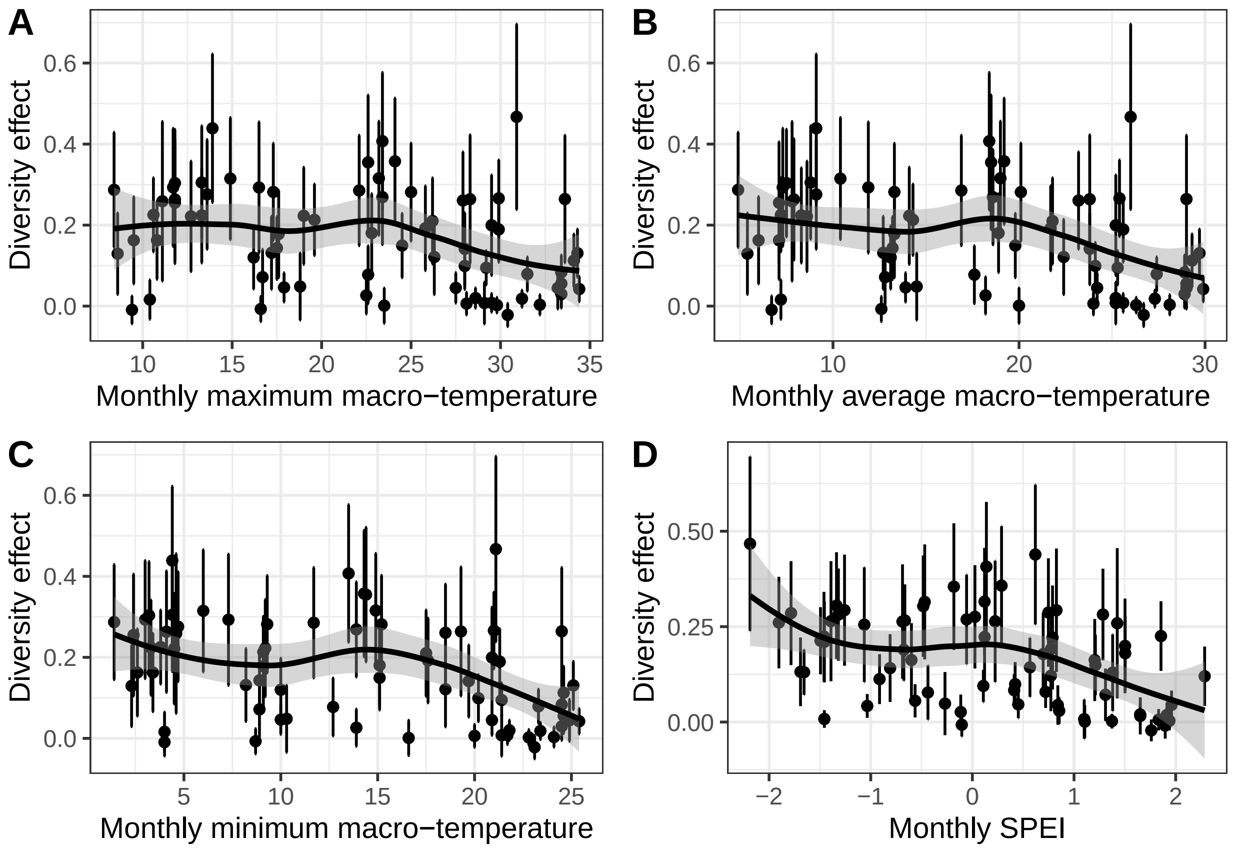

Figure S3 Macroclimate effects on the relationship between tree species richness and minimum microclimate temperature. Black points and error bars show diversity effects, i.e., slopes and respective standard errors, of the regression between minimum microclimate temperatures and tree species richness per month based on the model in Fig.1b (n = 63 for 12 months and 6 years). The change in these diversity effects with macroclimate conditions is examined for maximum (A), average (B) and minimum (C) monthly macroclimate temperatures and monthly values of the standardised precipitation evapotranspiration index (SPEI1; D). Trends are highlighted with a loess regression.

**Temperature buffering**

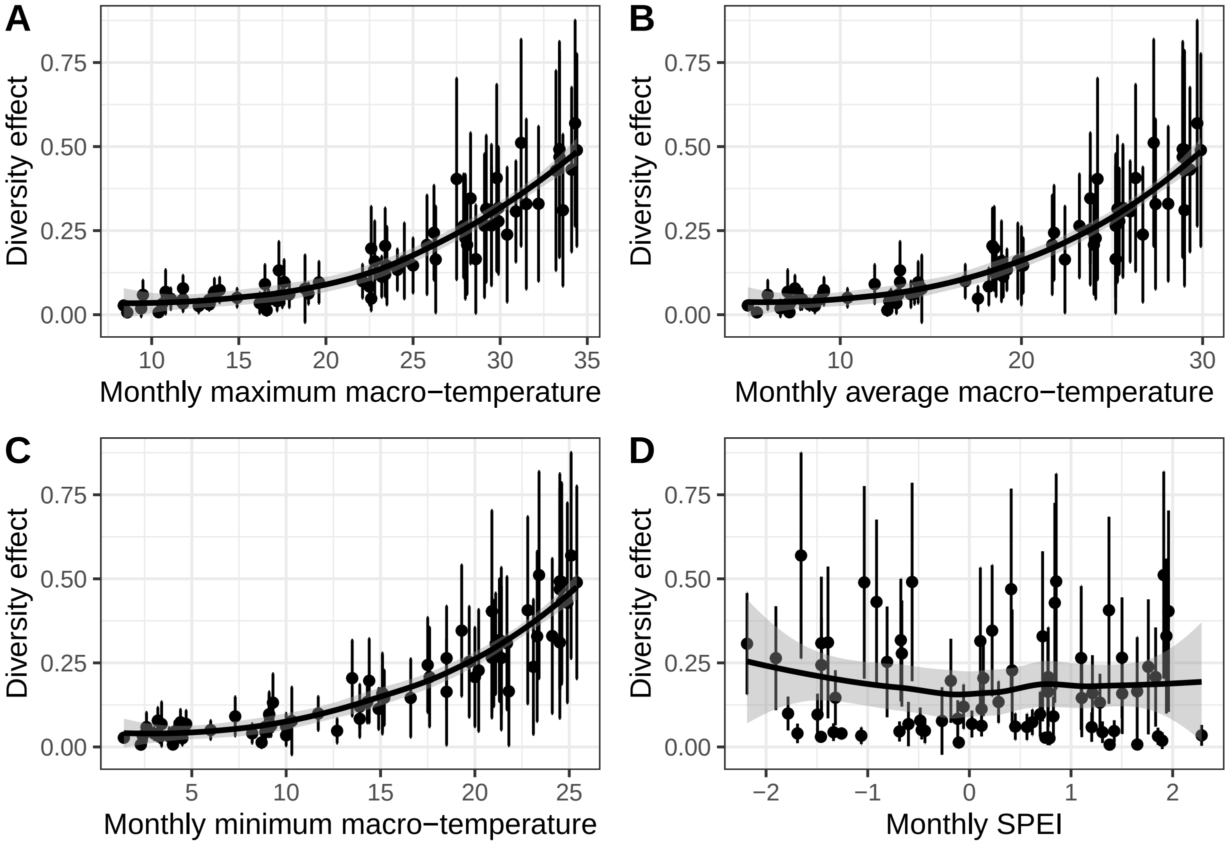

Figure S4 Macroclimate effects on the relationship between tree species richness and temperature buffering. Black points and error bars show diversity effects, i.e., slopes and respective standard errors, of the regression between temperature buffering and tree species richness per month based on the model in Fig.1b (n = 63 for 12 months and 6 years). The change in these diversity effects with macroclimate conditions is examined for maximum (A), average (B) and minimum (C) monthly macroclimate temperatures and monthly values of the standardised precipitation evapotranspiration index (SPEI1; D). Trends are highlighted with a loess regression.

### Supplementary S2

Florian Schnabel, Rémy Beugnon, Bo Yang, et al.

Tree diversity increases forest temperature buffering

#### Contents

Fig. 1 models ........ 2
Fig. 2 models ........ 10

#### Fig. 1

##### Fig. 1.A. Daily model

```
mod.daily =
  lme(T ~ log(TreeDiv) * hour.f,
      random = ~ 1|site/plot/date,
      data = data,
      correlation=corCAR1())
```

##### Model structure

###### Summary

| | Value | Std.Error | DF | t-value | p-value |
|---|---|---|---|---|---|
| (Intercept) | 16.461109580 | 0.29598888 | 699945 | 55.6139464 | 0.000000e+00 |
| log(TreeDiv) | 0.233754431 | 0.06120162 | 60 | 3.8194160 | 3.196367e-04 |
| hour.f1 | -0.353838995 | 0.01010693 | 699945 | -35.0095347 | 2.755591e-268 |
| hour.f2 | -0.671057326 | 0.01406768 | 699945 | -47.7020751 | 0.000000e+00 |
| hour.f3 | -0.954797225 | 0.01695454 | 699945 | -56.3151333 | 0.000000e+00 |
| hour.f4 | -1.192269321 | 0.01926166 | 699945 | -61.8985853 | 0.000000e+00 |
| hour.f5 | -1.408960925 | 0.02118343 | 699945 | -66.5123944 | 0.000000e+00 |
| hour.f6 | -1.474335370 | 0.02282088 | 699945 | -64.6046562 | 0.000000e+00 |
| hour.f7 | -0.770207530 | 0.02423462 | 699945 | -31.7812868 | 1.690983e-221 |
| hour.f8 | 1.113968491 | 0.02546441 | 699945 | 43.7461010 | 0.000000e+00 |
| hour.f9 | 3.622253242 | 0.02653790 | 699945 | 136.4936100 | 0.000000e+00 |
| hour.f10 | 5.915129984 | 0.02747515 | 699945 | 215.2901470 | 0.000000e+00 |
| hour.f11 | 7.497673735 | 0.02829174 | 699945 | 265.0128400 | 0.000000e+00 |
| hour.f12 | 8.452801951 | 0.02899930 | 699945 | 291.4829708 | 0.000000e+00 |
| hour.f13 | 8.865540613 | 0.02960540 | 699945 | 299.4568608 | 0.000000e+00 |
| hour.f14 | 8.861762002 | 0.03011680 | 699945 | 294.2464918 | 0.000000e+00 |
| hour.f15 | 8.260187516 | 0.03053863 | 699945 | 270.4832452 | 0.000000e+00 |
| hour.f16 | 6.959912636 | 0.03087471 | 699945 | 225.4243568 | 0.000000e+00 |
| hour.f17 | 4.875093606 | 0.03112774 | 699945 | 156.6157099 | 0.000000e+00 |
| hour.f18 | 2.597058096 | 0.03129941 | 699945 | 82.9746697 | 0.000000e+00 |
| hour.f19 | 1.050187680 | 0.03139049 | 699945 | 33.4556025 | 3.338612e-245 |
| hour.f20 | 0.198676082 | 0.03140088 | 699945 | 6.3270860 | 2.499842e-10 |
| hour.f21 | -0.439222499 | 0.03133058 | 699945 | -14.0189698 | 1.210087e-44 |
| hour.f22 | -0.953018736 | 0.03117739 | 699945 | -30.5676270 | 4.506214e-205 |
| hour.f23 | -1.372137166 | 0.03093834 | 699945 | -44.3507085 | 0.000000e+00 |
| log(TreeDiv):hour.f1 | -0.007143661 | 0.00931026 | 699945 | -0.7672892 | 4.429099e-01 |
| log(TreeDiv):hour.f2 | -0.014790095 | 0.01295845 | 699945 | -1.1413471 | 2.537259e-01 |
| log(TreeDiv):hour.f3 | -0.025269709 | 0.01561726 | 699945 | -1.6180628 | 1.056495e-01 |
| log(TreeDiv):hour.f4 | -0.038393480 | 0.01774189 | 699945 | -2.1640013 | 3.046459e-02 |
| log(TreeDiv):hour.f5 | -0.047875770 | 0.01951145 | 699945 | -2.4537267 | 1.413868e-02 |
| log(TreeDiv):hour.f6 | -0.079430902 | 0.02101899 | 699945 | -3.7790059 | 1.574687e-04 |
| log(TreeDiv):hour.f7 | -0.238110705 | 0.02232036 | 699945 | -10.6678696 | 1.445663e-26 |
| log(TreeDiv):hour.f8 | -0.557472710 | 0.02345217 | 699945 | -23.7706192 | 7.540890e-125 |
| log(TreeDiv):hour.f9 | -0.839211416 | 0.02443991 | 699945 | -34.3377388 | 3.528071e-258 |
| log(TreeDiv):hour.f10 | -0.960315656 | 0.02530206 | 699945 | -37.9540523 | 6.941835e-315 |
| log(TreeDiv):hour.f11 | -0.975222227 | 0.02605291 | 699945 | -37.4323695 | 2.344701e-306 |
| log(TreeDiv):hour.f12 | -1.017111978 | 0.02670322 | 699945 | -38.0894809 | 4.057498e-317 |
| log(TreeDiv):hour.f13 | -1.049806289 | 0.02725995 | 699945 | -38.5109354 | 0.000000e+00 |
| log(TreeDiv):hour.f14 | -0.995745313 | 0.02772931 | 699945 | -35.9094936 | 3.933989e-282 |
| log(TreeDiv):hour.f15 | -0.829690508 | 0.02811600 | 699945 | -29.5095448 | 2.848326e-191 |

```
log(TreeDiv):hour.f16 -0.617270705 0.02842355 699945 -21.7168736 1.539264e-104
log(TreeDiv):hour.f17 -0.351687831 0.02865441 699945 -12.2734287  1.268479e-34
log(TreeDiv):hour.f18 -0.162450362 0.02881011 699945  -5.6386586  1.714456e-08
log(TreeDiv):hour.f19 -0.064458253 0.02889134 699945  -2.2310578  2.567762e-02
log(TreeDiv):hour.f20 -0.043365917 0.02889799 699945  -1.5006552  1.334452e-01
log(TreeDiv):hour.f21 -0.040297824 0.02883002 699945  -1.3977731  1.621817e-01
log(TreeDiv):hour.f22 -0.047151700 0.02868541 699945  -1.6437519  1.002279e-01
log(TreeDiv):hour.f23 -0.054777525 0.02846121 699945  -1.9246380  5.427507e-02

Analysis of Deviance Table (Type II tests)

Response: T
                         Chisq Df Pr(>Chisq)
log(TreeDiv)             2.196  1     0.1384
hour.f              469459.826 23     <2e-16 ***
log(TreeDiv):hour.f   4751.744 23     <2e-16 ***
---
Signif. codes:  0 '***' 0.001 '**' 0.01 '*' 0.05 '.' 0.1 ' ' 1
```

##### Fig. 1.B. Monthly models

##### Maximum temperature

```
mod.monthly.Tmax =
  lme(T.max ~ log(TreeDiv, base = 2) * month.f,
      random = ~ 1|site/plot/year,
      data = data,
      correlation=corCAR1())
```

##### Model

##### Summary

```
                                      Value Std.Error   DF    t-value
(Intercept)                     19.57766903 0.5817134 4079 33.6551784
log(TreeDiv, base = 2)          -0.43824335 0.2537465   60 -1.7270913
month.f2                         4.36477422 0.1659925 4079 26.2950108
month.f3                         8.80419707 0.2042676 4079 43.1012849
month.f4                        14.05269258 0.2211456 4079 63.5449885
month.f5                        15.69114115 0.2294448 4079 68.3874360
month.f6                        16.86451156 0.2336064 4079 72.1919860
month.f7                        20.09719459 0.2357222 4079 85.2579557
month.f8                        20.45908892 0.2368049 4079 86.3964074
month.f9                        17.71960854 0.2374308 4079 74.6306136
month.f10                       14.24044749 0.2379099 4079 59.8564626
month.f11                        8.65274119 0.2380927 4079 36.3419023
month.f12                        1.33768140 0.2381159 4079  5.6177743
log(TreeDiv, base = 2):month.f2  -0.09123946 0.1054842 4079 -0.8649588
log(TreeDiv, base = 2):month.f3  -0.16797419 0.1298339 4079 -1.2937617
log(TreeDiv, base = 2):month.f4  -0.34357585 0.1406828 4079 -2.4422018
log(TreeDiv, base = 2):month.f5  -0.40972801 0.1459787 4079 -2.8067655
log(TreeDiv, base = 2):month.f6  -0.40007604 0.1486341 4079 -2.6916837
log(TreeDiv, base = 2):month.f7  -0.50955502 0.1499841 4079 -3.3973926
log(TreeDiv, base = 2):month.f8  -0.48352234 0.1506751 4079 -3.2090392
log(TreeDiv, base = 2):month.f9  -0.38887356 0.1510301 4079 -2.5748082
log(TreeDiv, base = 2):month.f10 -0.36869841 0.1512533 4079 -2.4376217
log(TreeDiv, base = 2):month.f11 -0.27510810 0.1513581 4079 -1.8175971
log(TreeDiv, base = 2):month.f12 -0.09992012 0.1514093 4079 -0.6599338
                                       p-value
(Intercept)                     2.384471e-219
log(TreeDiv, base = 2)           8.929651e-02
month.f2                        6.638462e-141
month.f3                         0.000000e+00
month.f4                         0.000000e+00
month.f5                         0.000000e+00
month.f6                         0.000000e+00
month.f7                         0.000000e+00
month.f8                         0.000000e+00
month.f9                         0.000000e+00
month.f10                        0.000000e+00
month.f11                       8.973263e-251
month.f12                        2.063279e-08
log(TreeDiv, base = 2):month.f2   3.871123e-01
```

```
log(TreeDiv, base = 2):month.f3   1.958210e-01
log(TreeDiv, base = 2):month.f4   1.464016e-02
log(TreeDiv, base = 2):month.f5   5.027915e-03
log(TreeDiv, base = 2):month.f6   7.138252e-03
log(TreeDiv, base = 2):month.f7   6.868255e-04
log(TreeDiv, base = 2):month.f8   1.342109e-03
log(TreeDiv, base = 2):month.f9   1.006450e-02
log(TreeDiv, base = 2):month.f10  1.482667e-02
log(TreeDiv, base = 2):month.f11  6.919907e-02
log(TreeDiv, base = 2):month.f12  5.093336e-01

Analysis of Deviance Table (Type II tests)

Response: T.max
                                 Chisq Df Pr(>Chisq)
log(TreeDiv, base = 2)             8.749  1   0.003098 **
month.f                        20081.560 11  < 2.2e-16 ***
log(TreeDiv, base = 2):month.f    18.491 11   0.070872 .
---
Signif. codes:  0 '***' 0.001 '**' 0.01 '*' 0.05 '.' 0.1 ' ' 1
```

##### Median temperature

```r
mod.monthly.Tmed =
  lme(T.med ~ log(TreeDiv, base = 2) * month.f,
      random = ~ 1|site/plot/year,
      data = data,
      correlation=corCAR1())
```

##### Model

##### Summary

```
                                      Value  Std.Error   DF      t-value
(Intercept)                     6.768830462 0.10226932 4079  66.18632746
log(TreeDiv, base = 2)         -0.006701877 0.04773377   60  -0.14040118
month.f2                        1.648061152 0.08921893 4079  18.47210234
month.f3                        5.721374668 0.09601214 4079  59.59011942
month.f4                       11.039676668 0.09684910 4079 113.98843328
month.f5                       15.214060502 0.09701029 4079 156.82934744
month.f6                       17.345918932 0.09703577 4079 178.75799540
month.f7                       19.771210385 0.09703979 4079 203.74332397
month.f8                       20.133788348 0.09704043 4079 207.47834547
month.f9                       16.825512296 0.09707864 4079 173.31838007
month.f10                      12.681436964 0.09717757 4079 130.49757351
month.f11                       7.514031087 0.09717914 4079  77.32144284
month.f12                       1.971003684 0.09714175 4079  20.28997574
log(TreeDiv, base = 2):month.f2  0.006319520 0.05669399 4079   0.11146719
log(TreeDiv, base = 2):month.f3  0.013335328 0.06101264 4079   0.21856663
log(TreeDiv, base = 2):month.f4  0.005368003 0.06162355 4079   0.08710961
log(TreeDiv, base = 2):month.f5 -0.011503335 0.06172640 4079  -0.18636006
log(TreeDiv, base = 2):month.f6 -0.008702268 0.06174265 4079  -0.14094418
log(TreeDiv, base = 2):month.f7  0.027324205 0.06174522 4079   0.44253149
log(TreeDiv, base = 2):month.f8  0.047749689 0.06174563 4079   0.77332905
log(TreeDiv, base = 2):month.f9  0.022842755 0.06174578 4079   0.36994842
log(TreeDiv, base = 2):month.f10 0.004725324 0.06176818 4079   0.07650094
log(TreeDiv, base = 2):month.f11 0.044895599 0.06176875 4079   0.72683359
log(TreeDiv, base = 2):month.f12 0.011412949 0.06176868 4079   0.18476919
                                    p-value
(Intercept)                    0.000000e+00
log(TreeDiv, base = 2)         8.888133e-01
month.f2                       3.114187e-73
month.f3                       0.000000e+00
month.f4                       0.000000e+00
month.f5                       0.000000e+00
month.f6                       0.000000e+00
month.f7                       0.000000e+00
month.f8                       0.000000e+00
month.f9                       0.000000e+00
month.f10                      0.000000e+00
month.f11                      0.000000e+00
month.f12                      2.802751e-87
log(TreeDiv, base = 2):month.f2 9.112514e-01
log(TreeDiv, base = 2):month.f3 8.269986e-01
log(TreeDiv, base = 2):month.f4 9.305887e-01
```

```
log(TreeDiv, base = 2):month.f5  8.521717e-01
log(TreeDiv, base = 2):month.f6  8.879210e-01
log(TreeDiv, base = 2):month.f7  6.581281e-01
log(TreeDiv, base = 2):month.f8  4.393725e-01
log(TreeDiv, base = 2):month.f9  7.114401e-01
log(TreeDiv, base = 2):month.f10 9.390243e-01
log(TreeDiv, base = 2):month.f11 4.673696e-01
log(TreeDiv, base = 2):month.f12 8.534193e-01

Analysis of Deviance Table (Type II tests)

Response: T.med
                                  Chisq Df Pr(>Chisq)
log(TreeDiv, base = 2)         7.7400e-02  1     0.7808
month.f                        1.5867e+05 11     <2e-16 ***
log(TreeDiv, base = 2):month.f 2.0134e+00 11     0.9984
---
Signif. codes:  0 '***' 0.001 '**' 0.01 '*' 0.05 '.' 0.1 ' ' 1
```

##### Minimum temperature

```r
mod.monthly.Tmin =
  lme(T.min ~ log(TreeDiv, base = 2) * month.f,
      random = ~ 1|site/plot/year,
      data = data,
      correlation=corCAR1())
```

##### Model

##### Summary

```
                                      Value  Std.Error   DF     t-value
(Intercept)                     -2.382067736 0.39538293 4079  -6.0247107
log(TreeDiv, base = 2)           0.230373627 0.07574149   60   3.0415776
month.f2                         0.651024277 0.14434421 4079   4.5102209
month.f3                         5.908516812 0.15379733 4079  38.4175518
month.f4                        10.122382156 0.15473265 4079  65.4185278
month.f5                        16.742223652 0.15489810 4079 108.0854018
month.f6                        21.178165737 0.15492048 4079 136.7034581
month.f7                        24.213576294 0.15492351 4079 156.2937480
month.f8                        23.817736338 0.15492392 4079 153.7382742
month.f9                        18.857430155 0.15498715 4079 121.6709273
month.f10                       12.938571364 0.15515341 4079  83.3921189
month.f11                        7.451452399 0.15515712 4079  48.0252030
month.f12                        0.944843400 0.15510123 4079   6.0917852
log(TreeDiv, base = 2):month.f2  0.007844573 0.09172322 4079   0.0855244
log(TreeDiv, base = 2):month.f3 -0.080967219 0.09773236 4079  -0.8284586
log(TreeDiv, base = 2):month.f4 -0.034794391 0.09844868 4079  -0.3534267
log(TreeDiv, base = 2):month.f5 -0.033229754 0.09855425 4079  -0.3371722
log(TreeDiv, base = 2):month.f6 -0.193409268 0.09856853 4079  -1.9621807
log(TreeDiv, base = 2):month.f7 -0.200310608 0.09857046 4079  -2.0321565
log(TreeDiv, base = 2):month.f8 -0.109700041 0.09857072 4079  -1.1129069
log(TreeDiv, base = 2):month.f9 -0.026438142 0.09857090 4079  -0.2682145
log(TreeDiv, base = 2):month.f10 0.015154855 0.09860853 4079   0.1536871
log(TreeDiv, base = 2):month.f11 -0.072531542 0.09860959 4079  -0.7355425
log(TreeDiv, base = 2):month.f12 0.017238217 0.09860954 4079   0.1748129
                                      p-value
(Intercept)                      1.843779e-09
log(TreeDiv, base = 2)           3.487242e-03
month.f2                         6.658050e-06
month.f3                        6.916878e-276
month.f4                         0.000000e+00
month.f5                         0.000000e+00
month.f6                         0.000000e+00
month.f7                         0.000000e+00
month.f8                         0.000000e+00
month.f9                         0.000000e+00
month.f10                        0.000000e+00
month.f11                        0.000000e+00
month.f12                        1.219753e-09
log(TreeDiv, base = 2):month.f2  9.318487e-01
log(TreeDiv, base = 2):month.f3  4.074593e-01
log(TreeDiv, base = 2):month.f4  7.237868e-01
```

```
log(TreeDiv, base = 2):month.f5   7.360044e-01
log(TreeDiv, base = 2):month.f6   4.980934e-02
log(TreeDiv, base = 2):month.f7   4.220248e-02
log(TreeDiv, base = 2):month.f8   2.658140e-01
log(TreeDiv, base = 2):month.f9   7.885478e-01
log(TreeDiv, base = 2):month.f10  8.778641e-01
log(TreeDiv, base = 2):month.f11  4.620515e-01
log(TreeDiv, base = 2):month.f12  8.612353e-01
```

```
Analysis of Deviance Table (Type II tests)

Response: T.min
                                  Chisq Df Pr(>Chisq)
log(TreeDiv, base = 2)           21.497  1  3.544e-06 ***
month.f                       96704.926 11  < 2.2e-16 ***
log(TreeDiv, base = 2):month.f   12.027 11     0.3616
---
Signif. codes:  0 '***' 0.001 '**' 0.01 '*' 0.05 '.' 0.1 ' ' 1
```

#### Fig. 2

##### Fig. 2.A. Monthly buffering

```
mod.monthly.buff =
  lme(T.buff ~ log(TreeDiv, base = 2) * month.f,
      random = ~ 1|site/plot/year,
      data = data,
      correlation=corCAR1())
```

##### Model

##### Summary

```
                                    Value  Std.Error   DF      t-value
(Intercept)                  1.4214005264 0.29537876 4079  4.812128388
log(TreeDiv, base = 2)       0.0394521811 0.05887598   60  0.670089630
month.f2                     0.0486057553 0.07200281 4079  0.675053591
month.f3                     0.7763554182 0.07322991 4079 10.601616114
month.f4                     1.5227744516 0.07373060 4079 20.653222066
month.f5                     3.1260400990 0.07403747 4079 42.222404194
month.f6                     4.4289297956 0.07419196 4079 59.695552433
month.f7                     4.4694149685 0.07426987 4079 60.178039077
month.f8                     4.3485705414 0.07430920 4079 58.519950921
month.f9                     3.3171168286 0.07435149 4079 44.613989788
month.f10                    2.2417452134 0.05249168 4079 42.706673014
month.f11                    1.1821582635 0.06430037 4079 18.384935949
month.f12                    0.2101235375 0.06946630 4079  3.024827156
log(TreeDiv, base = 2):month.f2  0.0003591565 0.04578383 4079  0.007844615
log(TreeDiv, base = 2):month.f3  0.0190240565 0.04656494 4079  0.408548919
log(TreeDiv, base = 2):month.f4  0.0684437810 0.04692937 4079  1.458442221
log(TreeDiv, base = 2):month.f5  0.2078084991 0.04712517 4079  4.409713337
log(TreeDiv, base = 2):month.f6  0.2742314467 0.04722379 4079  5.807060903
log(TreeDiv, base = 2):month.f7  0.3758246368 0.04727355 4079  7.949998808
log(TreeDiv, base = 2):month.f8  0.3931844490 0.04729866 4079  8.312802272
log(TreeDiv, base = 2):month.f9  0.2281642151 0.04731140 4079  4.822605096
log(TreeDiv, base = 2):month.f10 0.1236448692 0.03332447 4079  3.710332205
log(TreeDiv, base = 2):month.f11 0.0445162634 0.04086506 4079  1.089347851
log(TreeDiv, base = 2):month.f12 0.0177350919 0.04418430 4079  0.401388976
                                      p-value
(Intercept)                      1.547319e-06
log(TreeDiv, base = 2)           5.053716e-01
month.f2                         4.996800e-01
month.f3                         6.349728e-26
month.f4                         3.251098e-90
month.f5                        1.976263e-323
month.f6                         0.000000e+00
month.f7                         0.000000e+00
month.f8                         0.000000e+00
month.f9                         0.000000e+00
month.f10                        0.000000e+00
month.f11                        1.377963e-72
month.f12                        2.503255e-03
log(TreeDiv, base = 2):month.f2  9.937414e-01
log(TreeDiv, base = 2):month.f3  6.828922e-01
```

```
log(TreeDiv, base = 2):month.f4   1.447957e-01
log(TreeDiv, base = 2):month.f5   1.061742e-05
log(TreeDiv, base = 2):month.f6   6.842002e-09
log(TreeDiv, base = 2):month.f7   2.394777e-15
log(TreeDiv, base = 2):month.f8   1.258852e-16
log(TreeDiv, base = 2):month.f9   1.468635e-06
log(TreeDiv, base = 2):month.f10  2.097453e-04
log(TreeDiv, base = 2):month.f11  2.760649e-01
log(TreeDiv, base = 2):month.f12  6.881548e-01

Analysis of Deviance Table (Type II tests)

Response: T.buff
                                  Chisq Df Pr(>Chisq)
log(TreeDiv, base = 2)           12.655  1  0.0003746 ***
month.f                       14426.937 11  < 2.2e-16 ***
log(TreeDiv, base = 2):month.f  129.454 11  < 2.2e-16 ***
---
Signif. codes:  0 '***' 0.001 '**' 0.01 '*' 0.05 '.' 0.1 ' ' 1
```

##### Fig. 2.B. Yearly buffering

```
mod.yearly.buff =
  lme(T.buff ~ log(TreeDiv, base = 2) * year,
      random = ~ 1|site/plot,
      data = d.1.2,
      correlation=corCAR1(form = ~year))
```

##### Model

##### Summary

```
                                     Value   Std.Error  DF     t-value
(Intercept)                       2.0560786834 0.062738594 302  32.7721510
log(TreeDiv, base = 2)            0.0263776944 0.009466743  60   2.7863537
year2016                         -0.1025026980 0.011653328 302  -8.7960018
year2017                         -0.0674084330 0.012571246 302  -5.3621123
year2018                         -0.2010923146 0.012749915 302 -15.7720514
year2019                         -0.0306506353 0.012773274 302  -2.3995911
year2020                          0.0234895127 0.012833858 302   1.8302768
log(TreeDiv, base = 2):year2016 -0.0008995374 0.007452435 302  -0.1207038
log(TreeDiv, base = 2):year2017  0.0072923693 0.008039454 302   0.9070727
log(TreeDiv, base = 2):year2018  0.0020643885 0.008131625 302   0.2538716
log(TreeDiv, base = 2):year2019 -0.0046568272 0.008146604 302  -0.5716280
log(TreeDiv, base = 2):year2020  0.0080297196 0.008170093 302   0.9828186
                                      p-value
(Intercept)                      1.836480e-101
log(TreeDiv, base = 2)            7.125667e-03
year2016                          1.106657e-16
year2017                          1.637655e-07
year2018                          2.682032e-41
year2019                          1.701901e-02
year2020                          6.819402e-02
log(TreeDiv, base = 2):year2016   9.040059e-01
log(TreeDiv, base = 2):year2017   3.650915e-01
log(TreeDiv, base = 2):year2018   7.997675e-01
log(TreeDiv, base = 2):year2019   5.679992e-01
log(TreeDiv, base = 2):year2020   3.264834e-01
```

```
Analysis of Deviance Table (Type II tests)

Response: T.buff
                              Chisq Df Pr(>Chisq)
log(TreeDiv, base = 2)      12.7254  1  0.0003607 ***
year                       692.9748  5  < 2.2e-16 ***
log(TreeDiv, base = 2):year   4.3008  5  0.5069632
---
Signif. codes:  0 '***' 0.001 '**' 0.01 '*' 0.05 '.' 0.1 ' ' 1
```

##### SPEI model

```
Joining with `by = join_by(year)`
```

```
                                 Value   Std.Error  DF    t-value
(Intercept)                  1.9570887357 0.062472153 310 31.3273778
```

```
log(TreeDiv, base = 2)       0.0289820482 0.008395717  60  3.4520039
spei                         0.0556799665 0.006069463 310  9.1737876
log(TreeDiv, base = 2):spei -0.0007614258 0.003872263 310 -0.1966359
                                 p-value
(Intercept)                 4.586476e-98
log(TreeDiv, base = 2)      1.025641e-03
spei                        6.581195e-18
log(TreeDiv, base = 2):spei 8.442413e-01

Analysis of Deviance Table (Type II tests)

Response: T.buff
                               Chisq Df Pr(>Chisq)
log(TreeDiv, base = 2)       12.6422  1  0.0003771 ***
spei                        127.9597  1  < 2.2e-16 ***
log(TreeDiv, base = 2):spei   0.0387  1  0.8441125
---
Signif. codes:  0 '***' 0.001 '**' 0.01 '*' 0.05 '.' 0.1 ' ' 1
```

### Supplementary S3

Florian Schnabel, Rémy Beugnon, Bo Yang, et al.

Tree diversity increases forest temperature buffering

**Contents**

Hypothesis-driven SEM framework……………………………………………………2
Examined forest properties…………………………………………………………..3

We built a hypothesis-driven Structural Equation Model (SEM) framework to explain mechanisms behind observed tree species richness effects on temperature buffering (Fig. S5). This SEM framework was informed by prior knowledge of relationships between tree species richness and forest properties that in turn have been shown to affect temperature buffering (Table S1). However, the indirect effect of species richness via these forest properties on temperature buffering as well as their relative importance have been rarely assessed. We focussed on three potential pathways pertaining to tree canopy thickness, density and structural diversity (i.e. the variation of canopy elements) in three-dimensional space (Fig. S5). Based on literature-derived hypotheses (Table S1) we expected that tree species richness enhances canopy thickness, density and structural diversity (pathways 1, 3 and 5) and that these mediators in turn increase temperature buffering (pathways 2, 4 and 6). We additionally expected that species richness could potentially influence forest temperature buffering via processes not considered here (pathway 7).

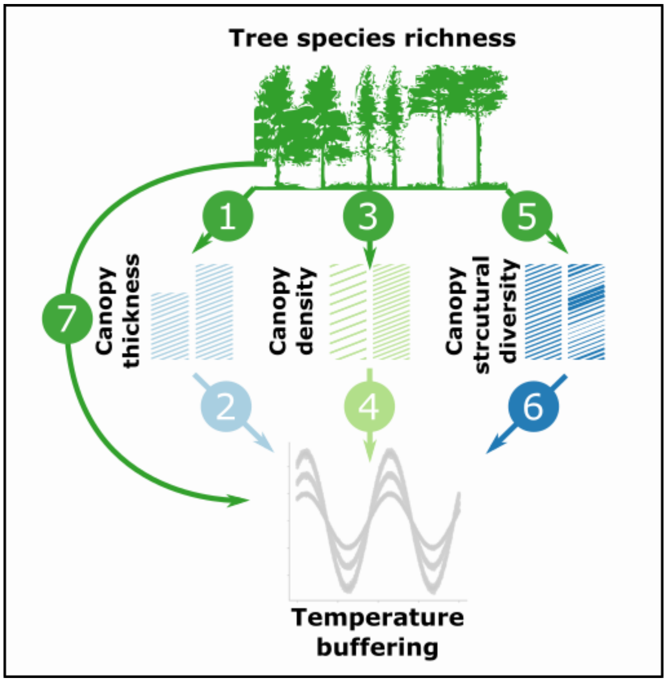

Figure S5 Hypothesis-driven SEM framework examining potential mediators of species richness effects on temperature buffering. Forest properties hypothesized to affect temperature buffering are show as sketches representing the thickness, density and structural diversity of aboveground canopy elements (e.g. leaves, branches or stems) at low (left) and high levels (right).

Table S1 Relationships between the variables considered in the SEM.

| Forest property | Pathway | Hypothesized mechanism |
|---|---|---|
| Canopy thickness | 1 | Tree diversity can increase mean stand-level tree height[1]. |
| | 2 | Mean stand-level tree height can increase temperature buffering[2,3]. |

| | | |
|---|---|---|
| Canopy density | 3 | Tree diversity can increase stand-level leaf area index (or basal area), through often-higher tree growth in mixtures[4–7]. |
| | 4 | Higher stand-level leaf area index (or basal area) can increase temperature buffering[3,8–11]. |
| Canopy structural diversity | 5 | Tree diversity can increase stand-level structural diversity, defined here as structural complexity in three-dimensional canopy space (SSCI index)[12,13]. |
| | 6 | Stand structural diversity can increase temperature buffering[2,14,15]. |
| Others | 7 | Species richness may also influence the temperature buffering of forests via other mechanisms not considered here. |

To quantify canopy thickness, density and structural diversity, we assembled a range of variables from former studies and tree inventories in the BEF-China experiment. Overall, we could capture plot-level (i.e. stand-level) canopy thickness through data on mean tree height, mean crown length and mean crown base height, canopy density through the summed basal area and leaf area index (LAI) and structural diversity through the terrestrial laser scanning (TLS) derived effective number of layers (ENL) and structural complexity index (SSCI)[14,16] (Table S2). Tree basal diameter (5cm above ground level), height and crown base height were measured for the central 6 × 6 trees in each VIP plot to avoid edge effects and subsequently used to derive plot-level means for the inventory-based forest properties. Peng et al. 2017 detail the methods used to quantify LAI[6] and Perles-Garcia et al. 2021[13] the methods used to measure ENL and SSCI. Of the potential variables (Table S2) we selected the ones with the highest relevance for temperature buffering according to literature-derived hypothesis (Table S1), basically focussing on the ones which were most successfully used as predictors of temperature buffering in former studies. Specifically, we selected mean tree height, LAI and SSCI as proxies for canopy thickness, density and structural diversity, respectively. We selected LAI over basal area as we considered LAI to more specifically capture canopy density.

Table S2 Forest properties. Potential variables describing tree canopy thickness, density and structural diversity available within the BEF-China experiment as well as their sample size, temporal extent and references for already published data. Selected variables for the SEMs highlighted in bold.

| Forest property | Data | Number of plots | Temporal extent | Reference |
|---|---|---|---|---|
| Canopy thickness | **Mean tree height** | **32** | **2019** | **Inventory; Unpublished data** |
| | Mean crown base height (i.e. height of lowest living branch) | 32 | 2019 | Inventory; Unpublished data |

| | | | | |
|---|---|---|---|---|
| | Mean crown length (i.e. tree height - crown base height) | 32 | 2019 | Inventory; Unpublished data |
| Canopy density | **Leaf Area Index (LAI)** | **54** | **2014** | **Peng et al 2017[6]** |
| | Summed basal area | 32 | 2019 | Inventory; Unpublished data |
| Canopy structural diversity | **Stand structural complexity index (SSCI)** based on TLS | **74** | **2019** | **Perles-Garcia et al. 2021** |
| | Effective number of layers (ENL) based on TLS | 74 | 2019 | Perles-Garcia et al. 2021[13] |

### Supplementary S4

Florian Schnabel, Rémy Beugnon, Bo Yang, et al.

Tree diversity increases forest temperature buffering

**Data**

Variable acronyms:

Buff: Monthly microclimate buffering (1/CV)

Buff.cor: Monthly microclimate buffering (1/CV) corrected by macroclimate

mean.height: mean stand height

l.SSCI: log-transformed SSCI index

LAI: Leaf Area Index

BA: Basal area

l.TreeDiv: log2-transformed tree species richness

**Correlation between variables**

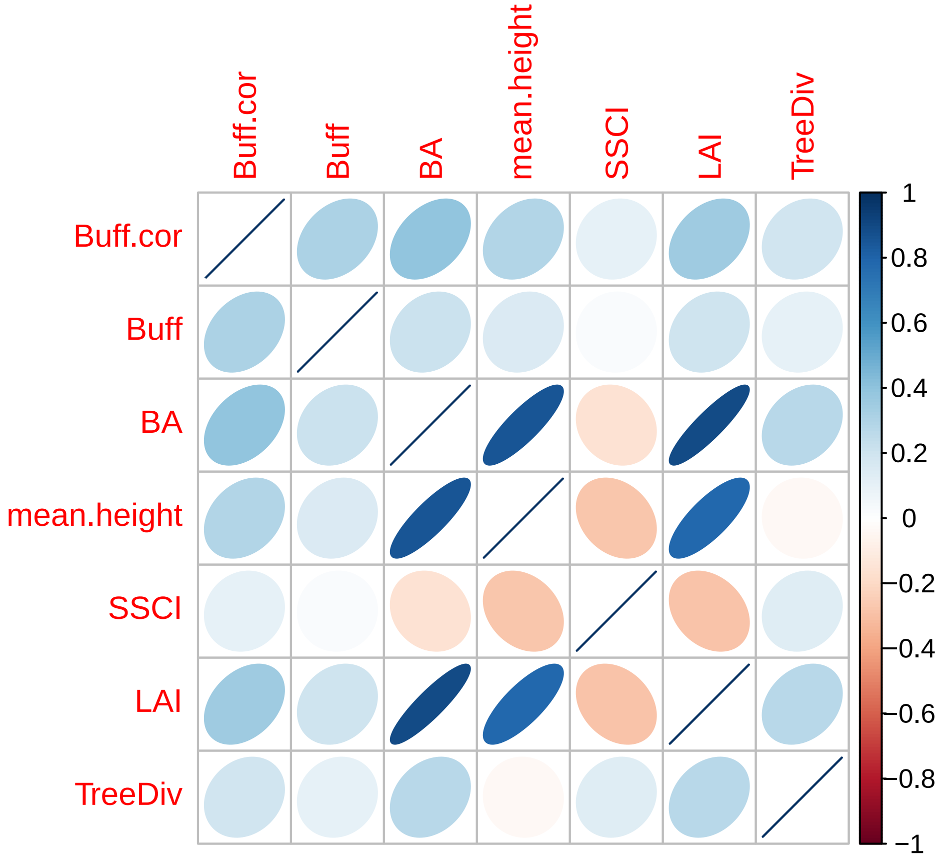

Figure S6 Correlation matrix between forest properties describing tree canopy thickness, density and structural diversity. Shown are Pearson's correlations between plot-level measurements of mean tree height (mean.height) as proxy for canopy thickness, basal area (BA) and leaf area index (LAI) as proxies for canopy density and structural complexity index (SSCI) as proxy for structural diversity. Correlations to monthly temperature buffering (Buff) and monthly temperature buffering controlled for macroclimate temperatures (Buff.cor) and tree species richness are included as well. Blue and red eclipses denote positive and negative correlations with thinner eclipses and darker colours showing stronger correlations; see the corrplot package in R for details.

**SEM at yearly scale**

```
Structural Equation Model of mod.sem

Call:
  Buff.cor ~ mean.height + l.SSCI + LAI + l.TreeDiv
  l.SSCI ~ l.TreeDiv
  LAI ~ l.TreeDiv + KoBi + ScSu + QuFa
  mean.height ~ l.TreeDiv
  LAI ~~ l.SSCI
  mean.height ~~ l.SSCI
  mean.height ~~ LAI

    AIC      BIC
 166.220   202.863

---
Tests of directed separation:

            Independ.Claim Test.Type DF Crit.Value P.Value
  mean.height ~ KoBi + ...      coef 24    -3.9027  0.0007 ***
       l.SSCI ~ KoBi + ...      coef 24     2.5928  0.0160   *
     Buff.cor ~ KoBi + ...      coef 21     1.8858  0.0732
  mean.height ~ ScSu + ...      coef 24     5.9524  0.0000 ***
       l.SSCI ~ ScSu + ...      coef 24     4.6599  0.0001 ***
     Buff.cor ~ ScSu + ...      coef 21     1.0154  0.3215
  mean.height ~ QuFa + ...      coef 24    -4.9291  0.0000 ***
       l.SSCI ~ QuFa + ...      coef 24     5.3072  0.0000 ***
     Buff.cor ~ QuFa + ...      coef 21    -0.4792  0.6367

Global goodness-of-fit:

  Fisher's C = 116.22 with P-value = 0 and on 18 degrees of freedom

---
Coefficients:

       Response   Predictor Estimate Std.Error DF Crit.Value P.Value
       Buff.cor mean.height   0.0705    0.1105 22     0.6383  0.5299
       Buff.cor      l.SSCI   0.2298     0.066 22     3.4833  0.0021
       Buff.cor         LAI   0.3284    0.1181 22     2.7805  0.0109
       Buff.cor   l.TreeDiv   0.0735     0.074 22     0.9941  0.3310
         l.SSCI   l.TreeDiv   0.1712    0.0551 72     3.1062  0.0027
            LAI   l.TreeDiv   0.8205    0.2915 49     2.8145  0.0070
            LAI        KoBi  -2.7238    0.8189 49    -3.3261  0.0017
            LAI        ScSu   3.1619    0.8475 49     3.7309  0.0005
            LAI        QuFa  -2.0357    0.9036 49    -2.2530  0.0288
    mean.height   l.TreeDiv  -3.0058   30.2082 30    -0.0995  0.9214
          ~~LAI    ~~l.SSCI   0.1140         - 74     0.9672  0.1684
  ~~mean.height    ~~l.SSCI  -0.1892         - 74    -1.6233  0.0545
  ~~mean.height       ~~LAI   0.2235         - 54     1.6379  0.0538
  Std.Estimate
        0.0705
        0.2298  **
```

```
       0.3284   *
       0.0735
       0.1712  **
       0.8205  **
      -0.9728  **
       1.1293 ***
      -0.7271   *
      -3.0058
       0.1140
      -0.1892
       0.2235

  Signif. codes:  0 '***' 0.001 '**' 0.01 '*' 0.05
```

```
---
Individual R-squared:

     Response method Marginal Conditional
     Buff.cor   none     0.19        0.91
       l.SSCI   none     0.12        0.89
          LAI   none     0.37        0.92
  mean.height   none     0.00        0.93
```

**Correlations addition to fullfill direct separation tests**

```
Structural Equation Model of mod.sem

Call:
  Buff.cor ~ mean.height + l.SSCI + LAI + l.TreeDiv
  l.SSCI ~ l.TreeDiv
  LAI ~ l.TreeDiv + KoBi + ScSu + QuFa
  mean.height ~ l.TreeDiv
  LAI ~~ l.SSCI
  mean.height ~~ l.SSCI
  mean.height ~~ LAI
  mean.height ~~ KoBi
  mean.height ~~ ScSu
  mean.height ~~ QuFa
  l.SSCI ~~ KoBi
  l.SSCI ~~ ScSu
  l.SSCI ~~ QuFa

    AIC      BIC
 58.401   95.044

---
Tests of directed separation:

         Independ.Claim Test.Type DF Crit.Value P.Value
  Buff.cor ~ KoBi + ...      coef 21     1.8858  0.0732
  Buff.cor ~ ScSu + ...      coef 21     1.0154  0.3215
  Buff.cor ~ QuFa + ...      coef 21    -0.4792  0.6367
```

```
Global goodness-of-fit:

  Fisher's C = 8.401 with P-value = 0.21 and on 6 degrees of freedom

---
Coefficients:

       Response   Predictor Estimate Std.Error  DF Crit.Value P.Value
       Buff.cor mean.height   0.0705    0.1105  22     0.6383  0.5299
       Buff.cor      l.SSCI   0.2298     0.066  22     3.4833  0.0021
       Buff.cor         LAI   0.3284    0.1181  22     2.7805  0.0109
       Buff.cor  l.TreeDiv    0.0735     0.074  22     0.9941   0.331
         l.SSCI  l.TreeDiv    0.1712    0.0551  72     3.1062  0.0027
            LAI  l.TreeDiv    0.8205    0.2915  49     2.8145   0.007
            LAI        KoBi  -2.7238    0.8189  49    -3.3261  0.0017
            LAI        ScSu   3.1619    0.8475  49     3.7309   5e-04
            LAI        QuFa  -2.0357    0.9036  49     -2.253  0.0288
    mean.height  l.TreeDiv   -3.0058   30.2082  30    -0.0995  0.9214
          ~~LAI    ~~l.SSCI    0.114         -  74     0.9672  0.1684
  ~~mean.height    ~~l.SSCI  -0.1892         -  74    -1.6233  0.0545
  ~~mean.height       ~~LAI   0.2235         -  54     1.6379  0.0538
  ~~mean.height      ~~KoBi        -         - 321          -       -
  ~~mean.height      ~~ScSu        -         - 321          -       -
  ~~mean.height      ~~QuFa  -0.0892         - 321    -1.5978  0.0555
       ~~l.SSCI      ~~KoBi    0.095         - 321     1.7026  0.0448
       ~~l.SSCI      ~~ScSu        -         - 321          -       -
       ~~l.SSCI      ~~QuFa  -0.1301         - 321      -2.34    0.01
  Std.Estimate
        0.0705
        0.2298  **
        0.3284   *
        0.0735
        0.1712  **
        0.8205  **
       -0.9728  **
        1.1293 ***
       -0.7271   *
       -3.0058
         0.114
       -0.1892
        0.2235
             -   -
             -   -
       -0.0892
         0.095   *
             -   -
       -0.1301  **

  Signif. codes:  0 '***' 0.001 '**' 0.01 '*' 0.05

---
Individual R-squared:

     Response method Marginal Conditional
```

```
     Buff.cor   none     0.19        0.91
       l.SSCI   none     0.12        0.89
          LAI   none     0.37        0.92
  mean.height   none     0.00        0.93
```

##### Monthly explicit models

```
[1] "Month =  1"
        Response   Predictor Estimate Std.Error DF Crit.Value P.Value
1       Buff.cor mean.height  -0.1555    0.2397 21    -0.6487  0.5235
2       Buff.cor      l.SSCI   0.2707    0.1424 21     1.9017  0.0710
3       Buff.cor         LAI   0.5181    0.2491 21     2.0799  0.0500
4       Buff.cor  l.TreeDiv  -0.0567    0.1496 21    -0.3787  0.7087
5         l.SSCI  l.TreeDiv   0.1587     0.068 33     2.3351  0.0258
6            LAI  l.TreeDiv   0.8128     0.334 26     2.4335  0.0221
7            LAI        KoBi  -2.5980    0.9164 26    -2.8351  0.0087
8            LAI        ScSu   3.1981    0.9627 26     3.3220  0.0027
9            LAI        QuFa  -2.0824    1.0041 26    -2.0739  0.0481
10   mean.height  l.TreeDiv  -3.0058   30.2082 30    -0.0995  0.9214
11         ~~LAI    ~~l.SSCI   0.0069         - 74     0.0577  0.4771
12 ~~mean.height    ~~l.SSCI  -0.1928         - 74    -1.6560  0.0511
13 ~~mean.height       ~~LAI   0.3036         - 54     2.2758  0.0135
14 ~~mean.height      ~~KoBi  -0.0764         - 32    -0.4127  0.3414
15 ~~mean.height      ~~ScSu  -0.1928         - 32    -1.0579  0.1494
16 ~~mean.height      ~~QuFa   0.0252         - 32     0.1357  0.4465
17      ~~l.SSCI      ~~KoBi  -0.1571         - 74    -1.3407  0.0921
18      ~~l.SSCI      ~~ScSu   0.3869         - 74     3.5351  0.0004
19      ~~l.SSCI      ~~QuFa   0.2423         - 74     2.1047  0.0194
   Std.Estimate
1       -0.2184
2        0.3489
3        0.7328   *
4       -0.0801
5        0.1741   *
6        0.8126   *
7       -0.9285  **
8        1.1430  **
9       -0.7442   *
10      -3.0260
11       0.0069
12      -0.1928
13       0.3036   *
14      -0.0764
15      -0.1928
16       0.0252
17      -0.1571
18       0.3869 ***
19       0.2423   *
  |                                                                          |
  |                                                                          |
[1] "Month =  2"
        Response   Predictor Estimate Std.Error DF Crit.Value P.Value
1       Buff.cor mean.height   0.0295    0.1202 21     0.2453  0.8086
```

```
2       Buff.cor      l.SSCI   0.1525    0.0714 21     2.1368  0.0445
3       Buff.cor         LAI   0.1584    0.1249 21     1.2689  0.2184
4       Buff.cor  l.TreeDiv  -0.0631     0.075 21    -0.8413  0.4096
5         l.SSCI  l.TreeDiv   0.1587     0.068 33     2.3351  0.0258
6            LAI  l.TreeDiv   0.8128     0.334 26     2.4335  0.0221
7            LAI        KoBi  -2.5980    0.9164 26    -2.8351  0.0087
8            LAI        ScSu   3.1981    0.9627 26     3.3220  0.0027
9            LAI        QuFa  -2.0824    1.0041 26    -2.0739  0.0481
10   mean.height  l.TreeDiv  -3.0058   30.2082 30    -0.0995  0.9214
11         ~~LAI    ~~l.SSCI   0.0069         - 74     0.0577  0.4771
12 ~~mean.height    ~~l.SSCI  -0.1928         - 74    -1.6560  0.0511
13 ~~mean.height       ~~LAI   0.3036         - 54     2.2758  0.0135
14 ~~mean.height      ~~KoBi  -0.0764         - 32    -0.4127  0.3414
15 ~~mean.height      ~~ScSu  -0.1928         - 32    -1.0579  0.1494
16 ~~mean.height      ~~QuFa   0.0252         - 32     0.1357  0.4465
17      ~~l.SSCI      ~~KoBi  -0.1571         - 74    -1.3407  0.0921
18      ~~l.SSCI      ~~ScSu   0.3869         - 74     3.5351  0.0004
19      ~~l.SSCI      ~~QuFa   0.2423         - 74     2.1047  0.0194
   Std.Estimate
1        0.0845
2        0.4011   *
3        0.4573
4       -0.1821
5        0.1741   *
6        0.8126   *
7       -0.9285  **
8        1.1430  **
9       -0.7442   *
10      -3.0260
11       0.0069
12      -0.1928
13       0.3036   *
14      -0.0764
15      -0.1928
16       0.0252
17      -0.1571
18       0.3869 ***
19       0.2423   *
  |                                                                          |
  |                                                                          |
[1] "Month =  3"
        Response   Predictor Estimate Std.Error DF Crit.Value P.Value
1       Buff.cor mean.height  -0.0973    0.0937 21    -1.0383  0.3110
2       Buff.cor      l.SSCI   0.1009    0.0556 21     1.8140  0.0840
3       Buff.cor         LAI   0.2430    0.0973 21     2.4963  0.0209
4       Buff.cor  l.TreeDiv  -0.0220    0.0585 21    -0.3762  0.7105
5         l.SSCI  l.TreeDiv   0.1587     0.068 33     2.3351  0.0258
6            LAI  l.TreeDiv   0.8128     0.334 26     2.4335  0.0221
7            LAI        KoBi  -2.5980    0.9164 26    -2.8351  0.0087
8            LAI        ScSu   3.1981    0.9627 26     3.3220  0.0027
9            LAI        QuFa  -2.0824    1.0041 26    -2.0739  0.0481
10   mean.height  l.TreeDiv  -3.0058   30.2082 30    -0.0995  0.9214
11         ~~LAI    ~~l.SSCI   0.0069         - 74     0.0577  0.4771
12 ~~mean.height    ~~l.SSCI  -0.1928         - 74    -1.6560  0.0511
```

```
13 ~~mean.height       ~~LAI   0.3036         - 54     2.2758  0.0135
14 ~~mean.height      ~~KoBi  -0.0764         - 32    -0.4127  0.3414
15 ~~mean.height      ~~ScSu  -0.1928         - 32    -1.0579  0.1494
16 ~~mean.height      ~~QuFa   0.0252         - 32     0.1357  0.4465
17      ~~l.SSCI      ~~KoBi  -0.1571         - 74    -1.3407  0.0921
18      ~~l.SSCI      ~~ScSu   0.3869         - 74     3.5351  0.0004
19      ~~l.SSCI      ~~QuFa   0.2423         - 74     2.1047  0.0194
   Std.Estimate
1       -0.3395
2        0.3232
3        0.8541   *
4       -0.0773
5        0.1741   *
6        0.8126   *
7       -0.9285  **
8        1.1430  **
9       -0.7442   *
10      -3.0260
11       0.0069
12      -0.1928
13       0.3036   *
14      -0.0764
15      -0.1928
16       0.0252
17      -0.1571
18       0.3869 ***
19       0.2423   *
  |                                                                          |
  |                                                                          |
[1] "Month =  4"
        Response   Predictor Estimate Std.Error DF Crit.Value P.Value
1       Buff.cor mean.height   0.0228    0.0673 22     0.3380  0.7386
2       Buff.cor      l.SSCI   0.1029    0.0402 22     2.5569  0.0180
3       Buff.cor         LAI   0.2096    0.0723 22     2.8989  0.0083
4       Buff.cor  l.TreeDiv   0.0097    0.0455 22     0.2131  0.8332
5         l.SSCI  l.TreeDiv   0.1587     0.068 33     2.3351  0.0258
6            LAI  l.TreeDiv   0.8128     0.334 26     2.4335  0.0221
7            LAI        KoBi  -2.5980    0.9164 26    -2.8351  0.0087
8            LAI        ScSu   3.1981    0.9627 26     3.3220  0.0027
9            LAI        QuFa  -2.0824    1.0041 26    -2.0739  0.0481
10   mean.height  l.TreeDiv  -3.0058   30.2082 30    -0.0995  0.9214
11         ~~LAI    ~~l.SSCI   0.0069         - 74     0.0577  0.4771
12 ~~mean.height    ~~l.SSCI  -0.1928         - 74    -1.6560  0.0511
13 ~~mean.height       ~~LAI   0.3036         - 54     2.2758  0.0135
14 ~~mean.height      ~~KoBi  -0.2541         - 32    -1.4148  0.0839
15 ~~mean.height      ~~ScSu   0.0416         - 32     0.2245  0.4120
16 ~~mean.height      ~~QuFa  -0.0381         - 32    -0.2055  0.4193
17      ~~l.SSCI      ~~KoBi   0.0167         - 74     0.1406  0.4443
18      ~~l.SSCI      ~~ScSu  -0.1942         - 74    -1.6680  0.0499
19      ~~l.SSCI      ~~QuFa  -0.1972         - 74    -1.6946  0.0473
   Std.Estimate
1        0.0772
2        0.3574  *
3        0.7091 **
```

```
4        0.0328
5        0.1546  *
6        0.8129  *
7       -0.9277 **
8        1.1421 **
9       -0.7436  *
10      -2.9994
11       0.0069
12      -0.1928
13       0.3036  *
14      -0.2541
15       0.0416
16      -0.0381
17       0.0167
18      -0.1942  *
19      -0.1972  *
  |                                                                          |
  |                                                                          |
[1] "Month =  5"
        Response   Predictor Estimate Std.Error DF Crit.Value P.Value
1       Buff.cor mean.height   0.1492    0.1436 22     1.0395  0.3099
2       Buff.cor      l.SSCI   0.1748    0.0858 22     2.0372  0.0538
3       Buff.cor         LAI   0.4034    0.1542 22     2.6170  0.0157
4       Buff.cor   l.TreeDiv   0.1499    0.0971 22     1.5441  0.1368
5         l.SSCI   l.TreeDiv   0.1587     0.068 33     2.3351  0.0258
6            LAI   l.TreeDiv   0.8128     0.334 26     2.4335  0.0221
7            LAI        KoBi  -2.5980    0.9164 26    -2.8351  0.0087
8            LAI        ScSu   3.1981    0.9627 26     3.3220  0.0027
9            LAI        QuFa  -2.0824    1.0041 26    -2.0739  0.0481
10   mean.height   l.TreeDiv  -3.0058   30.2082 30    -0.0995  0.9214
11         ~~LAI    ~~l.SSCI   0.0069         - 74     0.0577  0.4771
12 ~~mean.height    ~~l.SSCI  -0.1928         - 74    -1.6560  0.0511
13 ~~mean.height       ~~LAI   0.3036         - 54     2.2758  0.0135
14 ~~mean.height      ~~KoBi  -0.2541         - 32    -1.4148  0.0839
15 ~~mean.height      ~~ScSu   0.0416         - 32     0.2245  0.4120
16 ~~mean.height      ~~QuFa  -0.0381         - 32    -0.2055  0.4193
17      ~~l.SSCI      ~~KoBi   0.0167         - 74     0.1406  0.4443
18      ~~l.SSCI      ~~ScSu  -0.1942         - 74    -1.6680  0.0499
19      ~~l.SSCI      ~~QuFa  -0.1972         - 74    -1.6946  0.0473
   Std.Estimate
1        0.2145
2        0.2574
3        0.5787  *
4        0.2151
5        0.1546  *
6        0.8129  *
7       -0.9277 **
8        1.1421 **
9       -0.7436  *
10      -2.9994
11       0.0069
12      -0.1928
13       0.3036  *
14      -0.2541
```

```
15       0.0416
16      -0.0381
17       0.0167
18      -0.1942  *
19      -0.1972  *
  |                                                                            |
  |                                                                            |
[1] "Month =  6"
        Response   Predictor Estimate Std.Error DF Crit.Value P.Value
1       Buff.cor mean.height   0.1820    0.1993 22     0.9133  0.3710
2       Buff.cor      l.SSCI   0.2296    0.1191 22     1.9281  0.0668
3       Buff.cor         LAI   0.5510     0.214 22     2.5749  0.0173
4       Buff.cor   l.TreeDiv   0.1232    0.1348 22     0.9138  0.3707
5         l.SSCI   l.TreeDiv   0.1587     0.068 33     2.3351  0.0258
6            LAI   l.TreeDiv   0.8128     0.334 26     2.4335  0.0221
7            LAI        KoBi  -2.5980    0.9164 26    -2.8351  0.0087
8            LAI        ScSu   3.1981    0.9627 26     3.3220  0.0027
9            LAI        QuFa  -2.0824    1.0041 26    -2.0739  0.0481
10   mean.height   l.TreeDiv  -3.0058   30.2082 30    -0.0995  0.9214
11         ~~LAI    ~~l.SSCI   0.0069         - 74     0.0577  0.4771
12 ~~mean.height    ~~l.SSCI  -0.1928         - 74    -1.6560  0.0511
13 ~~mean.height       ~~LAI   0.3036         - 54     2.2758  0.0135
14 ~~mean.height      ~~KoBi  -0.2541         - 32    -1.4148  0.0839
15 ~~mean.height      ~~ScSu   0.0416         - 32     0.2245  0.4120
16 ~~mean.height      ~~QuFa  -0.0381         - 32    -0.2055  0.4193
17      ~~l.SSCI      ~~KoBi   0.0167         - 74     0.1406  0.4443
18      ~~l.SSCI      ~~ScSu  -0.1942         - 74    -1.6680  0.0499
19      ~~l.SSCI      ~~QuFa  -0.1972         - 74    -1.6946  0.0473
   Std.Estimate
1        0.2002
2        0.2587
3        0.6046  *
4        0.1352
5        0.1546  *
6        0.8129  *
7       -0.9277 **
8        1.1421 **
9       -0.7436  *
10      -2.9994
11       0.0069
12      -0.1928
13       0.3036  *
14      -0.2541
15       0.0416
16      -0.0381
17       0.0167
18      -0.1942  *
19      -0.1972  *
  |                                                                            |
  |                                                                            |
[1] "Month =  7"
        Response   Predictor Estimate Std.Error DF Crit.Value P.Value
1       Buff.cor mean.height   0.1598    0.1836 22     0.8704  0.3935
2       Buff.cor      l.SSCI   0.2975    0.1097 22     2.7112  0.0128
```

```
3       Buff.cor         LAI   0.5364    0.1971 22     2.7208  0.0125
4       Buff.cor  l.TreeDiv   0.0960    0.1242 22     0.7733  0.4476
5         l.SSCI  l.TreeDiv   0.1587     0.068 33     2.3351  0.0258
6            LAI  l.TreeDiv   0.8128     0.334 26     2.4335  0.0221
7            LAI        KoBi  -2.5980    0.9164 26    -2.8351  0.0087
8            LAI        ScSu   3.1981    0.9627 26     3.3220  0.0027
9            LAI        QuFa  -2.0824    1.0041 26    -2.0739  0.0481
10   mean.height  l.TreeDiv  -3.0058   30.2082 30    -0.0995  0.9214
11         ~~LAI    ~~l.SSCI   0.0069         - 74     0.0577  0.4771
12 ~~mean.height    ~~l.SSCI  -0.1928         - 74    -1.6560  0.0511
13 ~~mean.height       ~~LAI   0.3036         - 54     2.2758  0.0135
14 ~~mean.height      ~~KoBi  -0.2541         - 32    -1.4148  0.0839
15 ~~mean.height      ~~ScSu   0.0416         - 32     0.2245  0.4120
16 ~~mean.height      ~~QuFa  -0.0381         - 32    -0.2055  0.4193
17      ~~l.SSCI      ~~KoBi   0.0167         - 74     0.1406  0.4443
18      ~~l.SSCI      ~~ScSu  -0.1942         - 74    -1.6680  0.0499
19      ~~l.SSCI      ~~QuFa  -0.1972         - 74    -1.6946  0.0473
   Std.Estimate
1        0.1851
2        0.3530  *
3        0.6200  *
4        0.1110
5        0.1546  *
6        0.8129  *
7       -0.9277 **
8        1.1421 **
9       -0.7436  *
10      -2.9994
11       0.0069
12      -0.1928
13       0.3036  *
14      -0.2541
15       0.0416
16      -0.0381
17       0.0167
18      -0.1942  *
19      -0.1972  *
  |                                                                      |
  |                                                                      |
[1] "Month =  8"
        Response   Predictor Estimate Std.Error DF Crit.Value P.Value
1       Buff.cor mean.height   0.1517    0.1783 22     0.8510  0.4039
2       Buff.cor      l.SSCI   0.3353    0.1066 22     3.1465  0.0047
3       Buff.cor         LAI   0.6228    0.1915 22     3.2528  0.0036
4       Buff.cor  l.TreeDiv   0.1875    0.1206 22     1.5546  0.1343
5         l.SSCI  l.TreeDiv   0.1587     0.068 33     2.3351  0.0258
6            LAI  l.TreeDiv   0.8128     0.334 26     2.4335  0.0221
7            LAI        KoBi  -2.5980    0.9164 26    -2.8351  0.0087
8            LAI        ScSu   3.1981    0.9627 26     3.3220  0.0027
9            LAI        QuFa  -2.0824    1.0041 26    -2.0739  0.0481
10   mean.height  l.TreeDiv  -3.0058   30.2082 30    -0.0995  0.9214
11         ~~LAI    ~~l.SSCI   0.0069         - 74     0.0577  0.4771
12 ~~mean.height    ~~l.SSCI  -0.1928         - 74    -1.6560  0.0511
13 ~~mean.height       ~~LAI   0.3036         - 54     2.2758  0.0135
```

```
14 ~~mean.height      ~~KoBi  -0.2541         - 32    -1.4148  0.0839
15 ~~mean.height      ~~ScSu   0.0416         - 32     0.2245  0.4120
16 ~~mean.height      ~~QuFa  -0.0381         - 32    -0.2055  0.4193
17      ~~l.SSCI      ~~KoBi   0.0167         - 74     0.1406  0.4443
18      ~~l.SSCI      ~~ScSu  -0.1942         - 74    -1.6680  0.0499
19      ~~l.SSCI      ~~QuFa  -0.1972         - 74    -1.6946  0.0473
   Std.Estimate
1        0.1584
2        0.3585 **
3        0.6486 **
4        0.1953
5        0.1546  *
6        0.8129  *
7       -0.9277 **
8        1.1421 **
9       -0.7436  *
10      -2.9994
11       0.0069
12      -0.1928
13       0.3036  *
14      -0.2541
15       0.0416
16      -0.0381
17       0.0167
18      -0.1942  *
19      -0.1972  *
  |                                                                      |
  |                                                                      |
[1] "Month =  9"
        Response   Predictor Estimate Std.Error DF Crit.Value P.Value
1       Buff.cor mean.height   0.0675    0.1408 22     0.4790  0.6367
2       Buff.cor      l.SSCI   0.2678    0.0842 22     3.1821  0.0043
3       Buff.cor         LAI   0.3817    0.1512 22     2.5238  0.0193
4       Buff.cor   l.TreeDiv   0.1546    0.0953 22     1.6226  0.1189
5         l.SSCI   l.TreeDiv   0.1587     0.068 33     2.3351  0.0258
6            LAI   l.TreeDiv   0.8128     0.334 26     2.4335  0.0221
7            LAI        KoBi  -2.5980    0.9164 26    -2.8351  0.0087
8            LAI        ScSu   3.1981    0.9627 26     3.3220  0.0027
9            LAI        QuFa  -2.0824    1.0041 26    -2.0739  0.0481
10   mean.height   l.TreeDiv  -3.0058   30.2082 30    -0.0995  0.9214
11         ~~LAI    ~~l.SSCI   0.0069         - 74     0.0577  0.4771
12 ~~mean.height    ~~l.SSCI  -0.1928         - 74    -1.6560  0.0511
13 ~~mean.height       ~~LAI   0.3036         - 54     2.2758  0.0135
14 ~~mean.height      ~~KoBi  -0.2541         - 32    -1.4148  0.0839
15 ~~mean.height      ~~ScSu   0.0416         - 32     0.2245  0.4120
16 ~~mean.height      ~~QuFa  -0.0381         - 32    -0.2055  0.4193
17      ~~l.SSCI      ~~KoBi   0.0167         - 74     0.1406  0.4443
18      ~~l.SSCI      ~~ScSu  -0.1942         - 74    -1.6680  0.0499
19      ~~l.SSCI      ~~QuFa  -0.1972         - 74    -1.6946  0.0473
   Std.Estimate
1        0.1031
2        0.4192 **
3        0.5818  *
4        0.2357
```

```
5        0.1546  *
6        0.8129  *
7       -0.9277 **
8        1.1421 **
9       -0.7436  *
10      -2.9994
11       0.0069
12      -0.1928
13       0.3036  *
14      -0.2541
15       0.0416
16      -0.0381
17       0.0167
18      -0.1942  *
19      -0.1972  *
  |                                                                              |
  |                                                                              |
[1] "Month =  10"
        Response   Predictor Estimate Std.Error DF Crit.Value P.Value
1       Buff.cor mean.height   0.0537    0.0956 22     0.5618  0.5799
2       Buff.cor      l.SSCI   0.1883    0.0571 22     3.2976  0.0033
3       Buff.cor         LAI   0.1861    0.1026 22     1.8133  0.0834
4       Buff.cor   l.TreeDiv   0.0840    0.0646 22     1.2998  0.2071
5         l.SSCI   l.TreeDiv   0.1587     0.068 33     2.3351  0.0258
6            LAI   l.TreeDiv   0.8128     0.334 26     2.4335  0.0221
7            LAI        KoBi  -2.5980    0.9164 26    -2.8351  0.0087
8            LAI        ScSu   3.1981    0.9627 26     3.3220  0.0027
9            LAI        QuFa  -2.0824    1.0041 26    -2.0739  0.0481
10   mean.height   l.TreeDiv  -3.0058   30.2082 30    -0.0995  0.9214
11         ~~LAI    ~~l.SSCI   0.0069         - 74     0.0577  0.4771
12 ~~mean.height    ~~l.SSCI  -0.1928         - 74    -1.6560  0.0511
13 ~~mean.height       ~~LAI   0.3036         - 54     2.2758  0.0135
14 ~~mean.height      ~~KoBi  -0.2541         - 32    -1.4148  0.0839
15 ~~mean.height      ~~ScSu   0.0416         - 32     0.2245  0.4120
16 ~~mean.height      ~~QuFa  -0.0381         - 32    -0.2055  0.4193
17      ~~l.SSCI      ~~KoBi   0.0167         - 74     0.1406  0.4443
18      ~~l.SSCI      ~~ScSu  -0.1942         - 74    -1.6680  0.0499
19      ~~l.SSCI      ~~QuFa  -0.1972         - 74    -1.6946  0.0473
   Std.Estimate
1        0.1347
2        0.4841 **
3        0.4659
4        0.2104
5        0.1546  *
6        0.8129  *
7       -0.9277 **
8        1.1421 **
9       -0.7436  *
10      -2.9994
11       0.0069
12      -0.1928
13       0.3036  *
14      -0.2541
15       0.0416
```

```
16      -0.0381
17       0.0167
18      -0.1942  *
19      -0.1972  *
  |                                                                          |
  |                                                                          |
[1] "Month =  11"
        Response   Predictor Estimate Std.Error DF Crit.Value P.Value
1       Buff.cor mean.height  -0.0400    0.1212 22    -0.3302  0.7444
2       Buff.cor      l.SSCI   0.1980    0.0724 22     2.7333  0.0121
3       Buff.cor         LAI   0.2402    0.1301 22     1.8460  0.0784
4       Buff.cor   l.TreeDiv   0.0385     0.082 22     0.4696  0.6433
5         l.SSCI   l.TreeDiv   0.1587     0.068 33     2.3351  0.0258
6            LAI   l.TreeDiv   0.8128     0.334 26     2.4335  0.0221
7            LAI        KoBi  -2.5980    0.9164 26    -2.8351  0.0087
8            LAI        ScSu   3.1981    0.9627 26     3.3220  0.0027
9            LAI        QuFa  -2.0824    1.0041 26    -2.0739  0.0481
10   mean.height   l.TreeDiv  -3.0058   30.2082 30    -0.0995  0.9214
11         ~~LAI    ~~l.SSCI   0.0069         - 74     0.0577  0.4771
12 ~~mean.height    ~~l.SSCI  -0.1928         - 74    -1.6560  0.0511
13 ~~mean.height       ~~LAI   0.3036         - 54     2.2758  0.0135
14 ~~mean.height      ~~KoBi  -0.2541         - 32    -1.4148  0.0839
15 ~~mean.height      ~~ScSu   0.0416         - 32     0.2245  0.4120
16 ~~mean.height      ~~QuFa  -0.0381         - 32    -0.2055  0.4193
17      ~~l.SSCI      ~~KoBi   0.0167         - 74     0.1406  0.4443
18      ~~l.SSCI      ~~ScSu  -0.1942         - 74    -1.6680  0.0499
19      ~~l.SSCI      ~~QuFa  -0.1972         - 74    -1.6946  0.0473
   Std.Estimate
1       -0.0939
2        0.4758  *
3        0.5623
4        0.0901
5        0.1546  *
6        0.8129  *
7       -0.9277 **
8        1.1421 **
9       -0.7436  *
10      -2.9994
11       0.0069
12      -0.1928
13       0.3036  *
14      -0.2541
15       0.0416
16      -0.0381
17       0.0167
18      -0.1942  *
19      -0.1972  *
  |                                                                          |
  |                                                                          |
[1] "Month =  12"
        Response   Predictor Estimate Std.Error DF Crit.Value P.Value
1       Buff.cor mean.height  -0.0802    0.1245 22    -0.6444  0.5260
2       Buff.cor      l.SSCI   0.2065    0.0744 22     2.7742  0.0111
3       Buff.cor         LAI   0.2421    0.1337 22     1.8110  0.0838
```

```
4       Buff.cor   l.TreeDiv   0.0580    0.0842 22     0.6883  0.4985
5         l.SSCI   l.TreeDiv   0.1587     0.068 33     2.3351  0.0258
6            LAI   l.TreeDiv   0.8128     0.334 26     2.4335  0.0221
7            LAI        KoBi  -2.5980    0.9164 26    -2.8351  0.0087
8            LAI        ScSu   3.1981    0.9627 26     3.3220  0.0027
9            LAI        QuFa  -2.0824    1.0041 26    -2.0739  0.0481
10   mean.height   l.TreeDiv  -3.0058   30.2082 30    -0.0995  0.9214
11         ~~LAI    ~~l.SSCI   0.0069         - 74     0.0577  0.4771
12 ~~mean.height    ~~l.SSCI  -0.1928         - 74    -1.6560  0.0511
13 ~~mean.height       ~~LAI   0.3036         - 54     2.2758  0.0135
14 ~~mean.height      ~~KoBi  -0.2541         - 32    -1.4148  0.0839
15 ~~mean.height      ~~ScSu   0.0416         - 32     0.2245  0.4120
16 ~~mean.height      ~~QuFa  -0.0381         - 32    -0.2055  0.4193
17      ~~l.SSCI      ~~KoBi   0.0167         - 74     0.1406  0.4443
18      ~~l.SSCI      ~~ScSu  -0.1942         - 74    -1.6680  0.0499
19      ~~l.SSCI      ~~QuFa  -0.1972         - 74    -1.6946  0.0473
   Std.Estimate
1       -0.1845
2        0.4864  *
3        0.5556
4        0.1330
5        0.1546  *
6        0.8129  *
7       -0.9277 **
8        1.1421 **
9       -0.7436  *
10      -2.9994
11       0.0069
12      -0.1928
13       0.3036  *
14      -0.2541
15       0.0416
16      -0.0381
17       0.0167
18      -0.1942  *
19      -0.1972  *
  |                                                                         |
  |                                                                         |
```

| Month | Response | Predictor | Estimate | Std.Error | DF | P.Value | Std.Estimate | |
|---|---|---|---|---|---|---|---|---|
| 1 | Buff.cor | Mean height | -0.1555 | 0.2397 | 21 | 0.5235 | 0.0000 | |
| 1 | Buff.cor | SSCI | 0.2707 | 0.1424 | 21 | 0.0710 | 0.3489 | |
| 1 | Buff.cor | LAI | 0.5181 | 0.2491 | 21 | 0.0500 | 0.7328 | * |
| 1 | Buff.cor | Tree sp. richness | -0.0567 | 0.1496 | 21 | 0.7087 | 0.0000 | |
| 2 | Buff.cor | Mean height | 0.0295 | 0.1202 | 21 | 0.8086 | 0.0845 | |
| 2 | Buff.cor | SSCI | 0.1525 | 0.0714 | 21 | 0.0445 | 0.4011 | * |
| 2 | Buff.cor | LAI | 0.1584 | 0.1249 | 21 | 0.2184 | 0.4573 | |
| 2 | Buff.cor | Tree sp. richness | -0.0631 | 0.075 | 21 | 0.4096 | 0.0000 | |
| 3 | Buff.cor | Mean height | -0.0973 | 0.0937 | 21 | 0.3110 | 0.0000 | |
| 3 | Buff.cor | SSCI | 0.1009 | 0.0556 | 21 | 0.0840 | 0.3232 | |
| 3 | Buff.cor | LAI | 0.2430 | 0.0973 | 21 | 0.0209 | 0.8541 | * |
| 3 | Buff.cor | Tree sp. richness | -0.0220 | 0.0585 | 21 | 0.7105 | 0.0000 | |
| 4 | Buff.cor | Mean height | 0.0228 | 0.0673 | 22 | 0.7386 | 0.0772 | |
| 4 | Buff.cor | SSCI | 0.1029 | 0.0402 | 22 | 0.0180 | 0.3574 | * |
| 4 | Buff.cor | LAI | 0.2096 | 0.0723 | 22 | 0.0083 | 0.7091 | ** |
| 4 | Buff.cor | Tree sp. richness | 0.0097 | 0.0455 | 22 | 0.8332 | 0.0328 | |
| 5 | Buff.cor | Mean height | 0.1492 | 0.1436 | 22 | 0.3099 | 0.2145 | |
| 5 | Buff.cor | SSCI | 0.1748 | 0.0858 | 22 | 0.0538 | 0.2574 | |
| 5 | Buff.cor | LAI | 0.4034 | 0.1542 | 22 | 0.0157 | 0.5787 | * |
| 5 | Buff.cor | Tree sp. richness | 0.1499 | 0.0971 | 22 | 0.1368 | 0.2151 | |
| 6 | Buff.cor | Mean height | 0.1820 | 0.1993 | 22 | 0.3710 | 0.2002 | |
| 6 | Buff.cor | SSCI | 0.2296 | 0.1191 | 22 | 0.0668 | 0.2587 | |
| 6 | Buff.cor | LAI | 0.5510 | 0.214 | 22 | 0.0173 | 0.6046 | * |
| 6 | Buff.cor | Tree sp. richness | 0.1232 | 0.1348 | 22 | 0.3707 | 0.1352 | |
| 7 | Buff.cor | Mean height | 0.1598 | 0.1836 | 22 | 0.3935 | 0.1851 | |
| 7 | Buff.cor | SSCI | 0.2975 | 0.1097 | 22 | 0.0128 | 0.3530 | * |
| 7 | Buff.cor | LAI | 0.5364 | 0.1971 | 22 | 0.0125 | 0.6200 | * |
| 7 | Buff.cor | Tree sp. richness | 0.0960 | 0.1242 | 22 | 0.4476 | 0.1110 | |
| 8 | Buff.cor | Mean height | 0.1517 | 0.1783 | 22 | 0.4039 | 0.1584 | |
| 8 | Buff.cor | SSCI | 0.3353 | 0.1066 | 22 | 0.0047 | 0.3585 | ** |
| 8 | Buff.cor | LAI | 0.6228 | 0.1915 | 22 | 0.0036 | 0.6486 | ** |
| 8 | Buff.cor | Tree sp. richness | 0.1875 | 0.1206 | 22 | 0.1343 | 0.1953 | |
| 9 | Buff.cor | Mean height | 0.0675 | 0.1408 | 22 | 0.6367 | 0.1031 | |
| 9 | Buff.cor | SSCI | 0.2678 | 0.0842 | 22 | 0.0043 | 0.4192 | ** |
| 9 | Buff.cor | LAI | 0.3817 | 0.1512 | 22 | 0.0193 | 0.5818 | * |
| 9 | Buff.cor | Tree sp. richness | 0.1546 | 0.0953 | 22 | 0.1189 | 0.2357 | |
| 10 | Buff.cor | Mean height | 0.0537 | 0.0956 | 22 | 0.5799 | 0.1347 | |
| 10 | Buff.cor | SSCI | 0.1883 | 0.0571 | 22 | 0.0033 | 0.4841 | ** |
| 10 | Buff.cor | LAI | 0.1861 | 0.1026 | 22 | 0.0834 | 0.4659 | |
| 10 | Buff.cor | Tree sp. richness | 0.0840 | 0.0646 | 22 | 0.2071 | 0.2104 | |
| 11 | Buff.cor | Mean height | -0.0400 | 0.1212 | 22 | 0.7444 | 0.0000 | |
| 11 | Buff.cor | SSCI | 0.1980 | 0.0724 | 22 | 0.0121 | 0.4758 | * |
| 11 | Buff.cor | LAI | 0.2402 | 0.1301 | 22 | 0.0784 | 0.5623 | |
| 11 | Buff.cor | Tree sp. richness | 0.0385 | 0.082 | 22 | 0.6433 | 0.0901 | |
| 12 | Buff.cor | Mean height | -0.0802 | 0.1245 | 22 | 0.5260 | 0.0000 | |
| 12 | Buff.cor | SSCI | 0.2065 | 0.0744 | 22 | 0.0111 | 0.4864 | * |
| 12 | Buff.cor | LAI | 0.2421 | 0.1337 | 22 | 0.0838 | 0.5556 | |
| 12 | Buff.cor | Tree sp. richness | 0.0580 | 0.0842 | 22 | 0.4985 | 0.1330 | |

**SEM at yearly scale using Basal Area**

```
Structural Equation Model of mod.sem

Call:
  Buff.cor ~ mean.height + l.SSCI + BA + l.TreeDiv
  l.SSCI ~ l.TreeDiv
  BA ~ l.TreeDiv + KoBi + ScSu + QuFa
  mean.height ~ l.TreeDiv
  BA ~~ l.SSCI
  mean.height ~~ l.SSCI
  mean.height ~~ BA

    AIC      BIC
 171.937   210.046

---
Tests of directed separation:

            Independ.Claim Test.Type DF Crit.Value P.Value
  mean.height ~ KoBi + ...      coef 24    -3.9027  0.0007 ***
       l.SSCI ~ KoBi + ...      coef 24     2.5928  0.0160   *
     Buff.cor ~ KoBi + ...      coef 21     2.4195  0.0247   *
  mean.height ~ ScSu + ...      coef 24     5.9524  0.0000 ***
       l.SSCI ~ ScSu + ...      coef 24     4.6599  0.0001 ***
     Buff.cor ~ ScSu + ...      coef 21     1.6730  0.1092
  mean.height ~ QuFa + ...      coef 24    -4.9291  0.0000 ***
       l.SSCI ~ QuFa + ...      coef 24     5.3072  0.0000 ***
     Buff.cor ~ QuFa + ...      coef 21    -0.1696  0.8670

Global goodness-of-fit:

  Fisher's C = 119.937 with P-value = 0 and on 18 degrees of freedom

---
Coefficients:

       Response   Predictor Estimate Std.Error  DF Crit.Value P.Value
       Buff.cor mean.height  -0.0325    0.1418  22    -0.2289  0.8210
       Buff.cor      l.SSCI   0.1628    0.0638  22     2.5535  0.0181
       Buff.cor          BA   0.4093    0.1463  22     2.7974  0.0105
       Buff.cor   l.TreeDiv   0.0609    0.0762  22     0.7991  0.4328
         l.SSCI   l.TreeDiv   0.1712    0.0551  72     3.1062  0.0027
             BA   l.TreeDiv   0.2267    0.1098  27     2.0648  0.0487
             BA        KoBi  -0.9191    0.3641  27    -2.5242  0.0178
             BA        ScSu   1.4978    0.4021  27     3.7253  0.0009
             BA        QuFa  -0.8922    0.4021  27    -2.2191  0.0351
    mean.height   l.TreeDiv  -3.0058   30.2082  30    -0.0995  0.9214
           ~~BA    ~~l.SSCI   0.0003         - 192     0.0047  0.4981
  ~~mean.height    ~~l.SSCI  -0.1892         -  74    -1.6233  0.0545
  ~~mean.height        ~~BA  -0.0158         - 192    -0.2175  0.4140
  Std.Estimate
      -0.0325
       0.1628   *
```

```
        0.4093   *
        0.0609
        0.1712  **
        0.2267   *
       -0.3283   *
        0.5350 ***
       -0.3187   *
       -3.0058
        0.0003
       -0.1892
       -0.0158

  Signif. codes:  0 '***' 0.001 '**' 0.01 '*' 0.05

---
Individual R-squared:

     Response method Marginal Conditional
     Buff.cor   none     0.19        0.91
       l.SSCI   none     0.12        0.89
           BA   none     0.32        0.71
  mean.height   none     0.00        0.93
```

Correlation addition to fulfill the direct separation tests

```
Structural Equation Model of mod.sem

Call:
  Buff.cor ~ mean.height + l.SSCI + BA + l.TreeDiv
  l.SSCI ~ l.TreeDiv
  BA ~ l.TreeDiv + KoBi + ScSu + QuFa
  mean.height ~ l.TreeDiv
  BA ~~ l.SSCI
  mean.height ~~ l.SSCI
  mean.height ~~ BA
  mean.height ~~ KoBi
  mean.height ~~ ScSu
  mean.height ~~ QuFa
  l.SSCI ~~ QuFa
  l.SSCI ~~ ScSu

    AIC      BIC
 72.393   110.502

---
Tests of directed separation:

         Independ.Claim Test.Type DF Crit.Value P.Value
    l.SSCI ~ KoBi + ...      coef 24     2.5928  0.0160 *
  Buff.cor ~ KoBi + ...      coef 21     2.4195  0.0247 *
  Buff.cor ~ ScSu + ...      coef 21     1.6730  0.1092
  Buff.cor ~ QuFa + ...      coef 21    -0.1696  0.8670

Global goodness-of-fit:
```

```
  Fisher's C = 20.393 with P-value = 0.009 and on 8 degrees of freedom

---
Coefficients:

       Response   Predictor Estimate Std.Error  DF Crit.Value P.Value
       Buff.cor mean.height  -0.0325    0.1418  22    -0.2289   0.821
       Buff.cor      l.SSCI   0.1628    0.0638  22     2.5535  0.0181
       Buff.cor          BA   0.4093    0.1463  22     2.7974  0.0105
       Buff.cor  l.TreeDiv   0.0609    0.0762  22     0.7991  0.4328
         l.SSCI  l.TreeDiv   0.1712    0.0551  72     3.1062  0.0027
             BA  l.TreeDiv   0.2267    0.1098  27     2.0648  0.0487
             BA        KoBi  -0.9191    0.3641  27    -2.5242  0.0178
             BA        ScSu   1.4978    0.4021  27     3.7253   9e-04
             BA        QuFa  -0.8922    0.4021  27    -2.2191  0.0351
    mean.height  l.TreeDiv  -3.0058   30.2082  30    -0.0995  0.9214
           ~~BA    ~~l.SSCI    3e-04         - 192     0.0047  0.4981
  ~~mean.height    ~~l.SSCI  -0.1892         -  74    -1.6233  0.0545
  ~~mean.height        ~~BA  -0.0158         - 192    -0.2175   0.414
  ~~mean.height      ~~KoBi        -         - 321          -       -
  ~~mean.height      ~~ScSu        -         - 321          -       -
  ~~mean.height      ~~QuFa  -0.0892         - 321    -1.5978  0.0555
       ~~l.SSCI      ~~QuFa  -0.1301         - 321      -2.34    0.01
       ~~l.SSCI      ~~ScSu        -         - 321          -       -
  Std.Estimate
       -0.0325
        0.1628   *
        0.4093   *
        0.0609
        0.1712  **
        0.2267   *
       -0.3283   *
         0.535 ***
       -0.3187   *
       -3.0058
         3e-04
       -0.1892
       -0.0158
             -   -
             -   -
       -0.0892
       -0.1301  **
             -   -

  Signif. codes:  0 '***' 0.001 '**' 0.01 '*' 0.05

---
Individual R-squared:

     Response method Marginal Conditional
     Buff.cor   none     0.19        0.91
       l.SSCI   none     0.12        0.89
           BA   none     0.32        0.71
```

| | | | |
|---|---|---|---|
| mean.height | none | 0.00 | 0.93 |

### Supplementary S5

Florian Schnabel, Rémy Beugnon, Bo Yang, et al.

Tree diversity increases forest temperature buffering

**Diversity treatment** **5**

#### Spatial cover

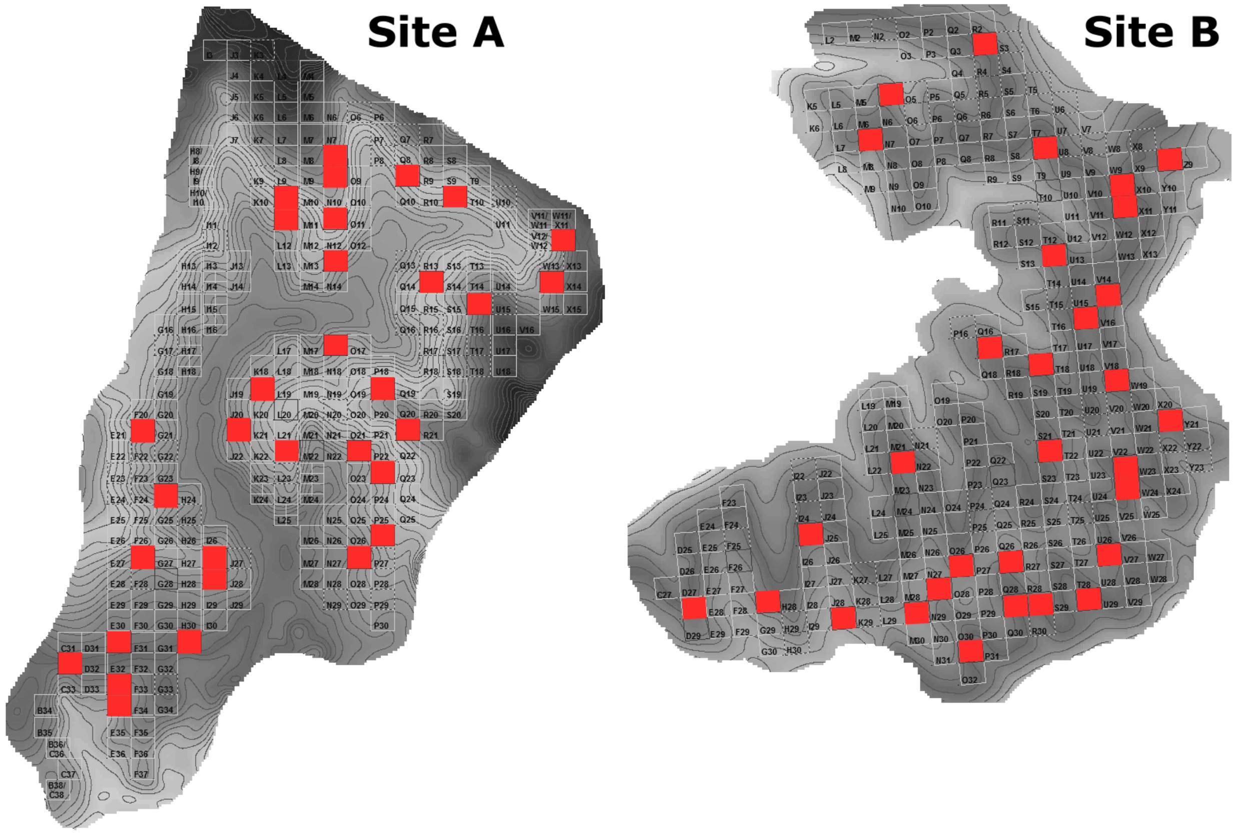

Figure S7 Position of the VIP plots within site A (n = 32) and B (n = 31) of the BEF-China experiment which are equipped with temperature loggers. Note, one plot at site B was excluded due to logger malfunction.

### Temporal structure

#### Temporal cover

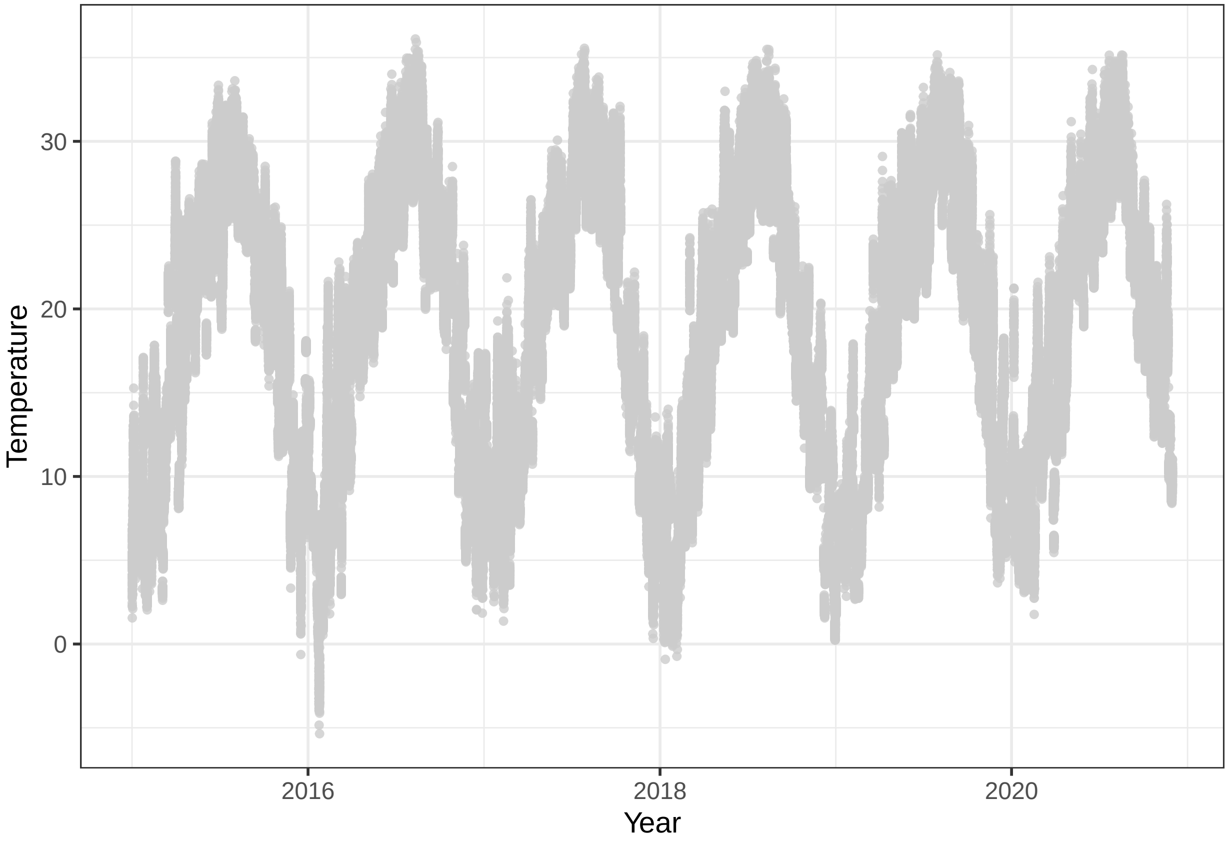

Figure S8 Time series of microclimate temperature measurements across years to illustrate the annual course of microclimate temperatures. Grey points shown hourly temperature measurements.

#### Temporal resolution

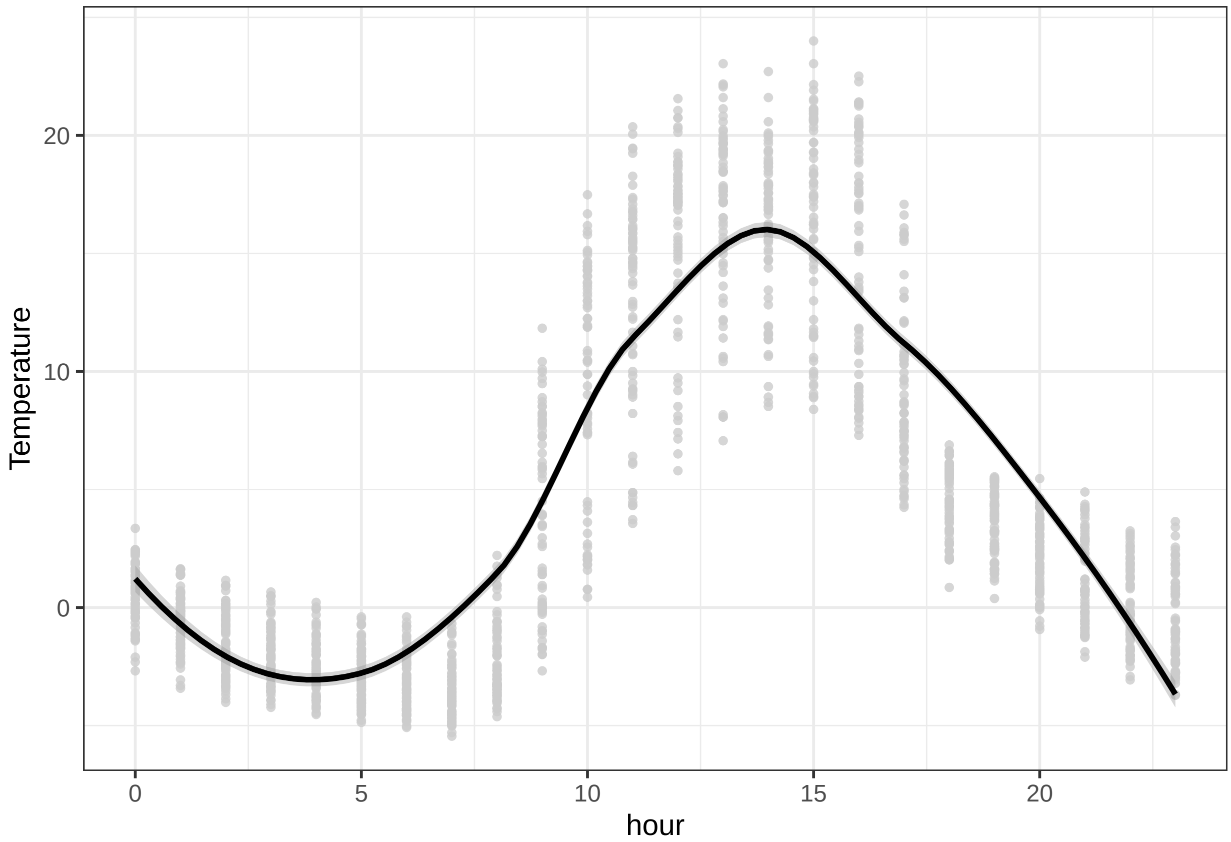

Figure S9 Resolution of the time series hourly measurement (here 01.01.2015) to illustrate the daily course of microclimate temperatures. Grey points shown hourly temperature measurements for all plots.

#### Diversity treatment

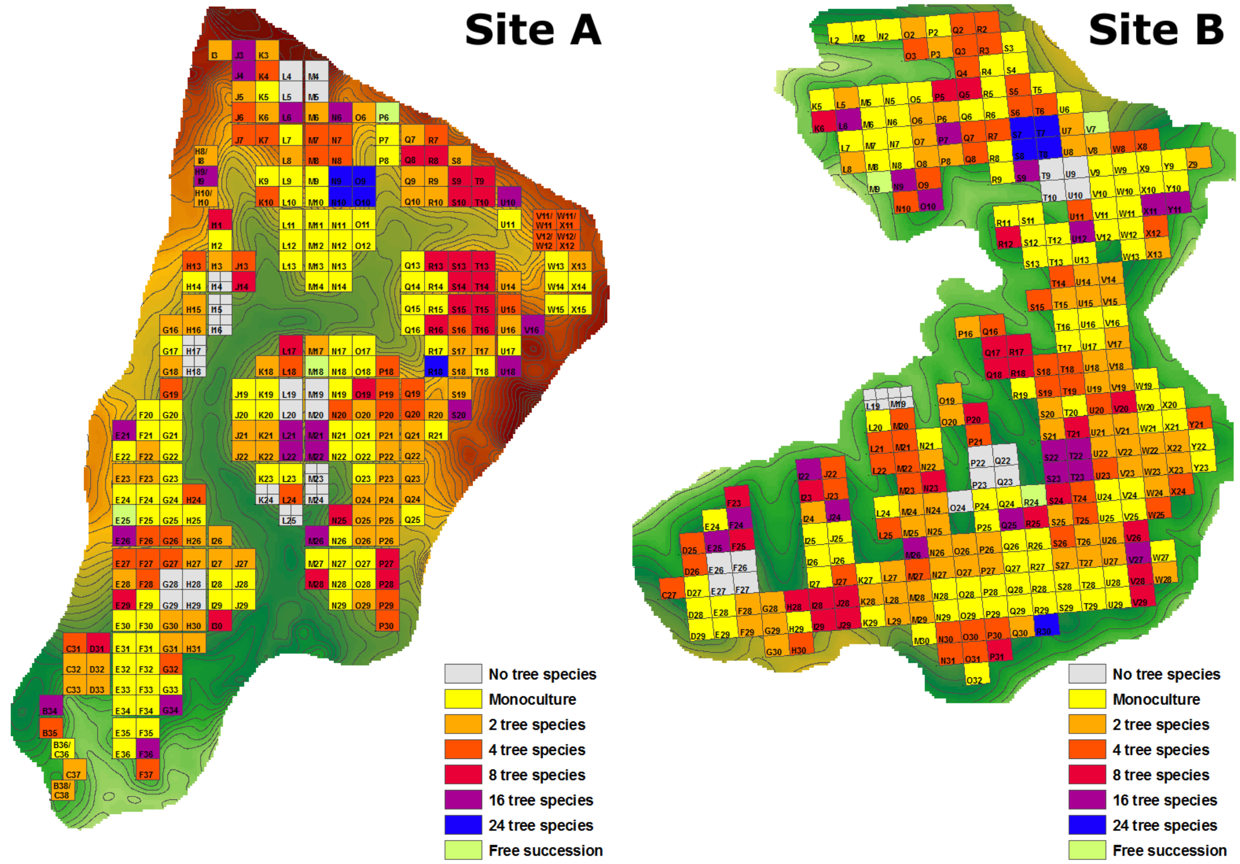

Figure S10 Position of plots covering a species richness gradient of 1 to 24 tree species within site A and B of the BEF-China experiment. We examined 32 VIP plots at site A and 31 VIP plots at site B covering the entire species richness gradient (the number of plots per richness level is shown in Table S3).

| Tree species richness | Number of plots |
|---|---|
| 1 | 32 |
| 2 | 16 |
| 4 | 7 |
| 8 | 4 |
| 16 | 2 |
| 24 | 2 |

Table S3 Tree diversity experimental design
